# Dynamic regulation of MBNL1 dosage through a stress-responsive intronic polyadenylation pathway

**DOI:** 10.64898/2026.11.26.769111

**Authors:** Katarzyna Taylor, Marta M. Jakimowicz, Klaudia Kubiak, Daniela C. Hernandez Enseñat, Anna Baud, Eliza Kopera, Magdalena M. Masłoń, Łukasz J. Sznajder, Krzysztof Sobczak

**Affiliations:** Department of Gene Expression, Institute of Molecular Biology and Biotechnology, Adam Mickiewicz University, Poznań, Poland; Department of Chemistry and Biochemistry, University of Nevada, Las Vegas, NV, USA; Center for Development of Gene Therapies, Center for Advanced Technologies, Adam Mickiewicz University, Poznań, Poland; Małopolska Centre of Biotechnology, Jagiellonian University, Kraków, Poland

**Keywords:** MBNL, intronic polyadenylation, myotonic dystrophy, cellular stress

## Abstract

Muscleblind like splicing regulator 1 (MBNL1) is a key RNA-binding protein that orchestrates a developmental splicing program during cell maturation and differentiation across multiple tissues, including skeletal and cardiac muscle, the central nervous system, and the immune system. Dysregulation of MBNL1 contributes to the pathogenesis of cancer and tandem repeat expansion disorders, including myotonic dystrophy type 1 (DM1). Although MBNL1 dosage is a major determinant of the MBNL1-dependent RNA processing program, the mechanisms governing *MBNL1* expression remain incompletely understood. Here, we identify an intronic polyadenylation (IPA) pathway within the *MBNL1* locus as a novel, post-transcriptional regulatory mechanism. We show that an intronic polyadenylation site within the exceptionally long intron 1 of *MBNL1*, coupled with splicing and exonization of an alternative last exon (ALE), generates a stable polyadenylated transcript, termed *MBNL1-IPA*, that attenuates the expression of canonical *MBNL1* isoforms. *MBNL1-IPA* is enriched in immune cells and is actively suppressed in non-immune cells through combined effects of transcription kinetics, splicing factor proline and glutamine (SFPQ)-dependent maintenance of long-intron integrity, and a local *cis*-regulatory element. Disruption of any of these regulatory layers promotes ALE inclusion and reduces canonical *MBNL1* expression. We further demonstrate that innate immune stress activates the IPA pathway, leading to reduced MBNL1 protein abundance across various cell types and altered MBNL1-dependent RNA processing in DM1-derived fibroblasts. Conversely, CRISPR-Cas9-mediated deletion of the ALE region increases *MBNL1* expression under basal conditions and further enhances its expression during innate immune stress. Together, our findings establish the *MBNL1* IPA pathway as a stress-responsive mechanism that fine-tunes MBNL1 dosage and reveal a previously unappreciated regulatory layer with potential relevance to the pathogenesis and therapeutic modulation of DM1 and other disorders associated with MBNL1 dysfunction.

## INTRODUCTION

Precise regulation of RNA processing is essential for maintaining cellular identity and adaptation to changing physiological conditions. Among post-transcriptional mechanisms, alternative splicing (AS) and alternative polyadenylation (APA) including IPA play central roles in shaping transcript output and proteome diversity^1–3^. These processes are coordinated by RNA-binding proteins (RBPs), which frequently participate in autoregulatory and cross-regulatory networks that maintain RNA-processing homeostasis^4–6^. IPA has recently emerged as an important mechanism regulating output of long genes at the level of RNA processing^7–9^. Cleavage and polyadenylation within introns can attenuate the production of full-length transcripts^10^ and sometimes generate stable alternative non-coding RNAs or micropeptides^11–14^. In mammalian cells, IPA is particularly influenced by intron architecture, transcription elongation kinetics, and the activity of RBPs and splicing factors^8–10, 15^. Importantly, IPA can be selectively engaged under specific physiological conditions, including immune activation, neuronal differentiation, stress responses, and cancer-associated states, where it contributes to rapid remodeling of transcript output^7, 16–21^. Whether similar long intron-associated mechanisms participate in regulating the expression of dosage-sensitive splicing factors remains largely unexplored.

Muscleblind-like (MBNL) splicing regulators belong to a family of RBPs that are involved in several critical aspects of RNA metabolism during prenatal and postnatal development. These include regulation of AS and APA programs, mRNA stability, RNA transport and localization as well as circular RNA and microRNA biogenesis^22–31^. The MBNL1 and MBNL2 paralogs are essential for proper development and homeostasis of various tissues; however, MBNL1 is a major regulator in skeletal muscles^30, 32^ and the immune system^24, 33^, whereas MBNL2 contributes primarily to brain and heart function^34–36^. In contrast, MBNL3 function is largely restricted to embryonic stages and tissue regeneration^37, 38^. Notably, MBNL-dependent splicing events are differentially sensitive to MBNL dosage, such that even partial reduction or increase in MBNL activity can result in significant alterations in the downstream splicing program^39, 40^.

Aberrant *MBNL* expression and the resulting disruption of MBNL-dependent RNA processing are hallmarks of several cancers and tandem repeat expansion disorders^27^. Depending on the cellular context, MBNL1 can function either as an oncogene^41, 42^ or as a tumor suppressor and has emerged as an important regulator of the tumor immune microenvironment^43^. Its oncogenic role has been particularly well characterized in rearranged leukemias, where MBNL1 has been proposed as a potential therapeutic target^42, 44, 45^. Conversely, functional depletion of MBNL proteins underlies the pathogenesis of several genetic disorders, including Fuchs’ endothelial corneal dystrophy (FECD) and myotonic dystrophy type 1 (DM1) and type 2 (DM2)^46^. In FECD, DM1 and DM2, expanded C/CUG repeat-containing transcripts produced from the *TCF4*, *DMPK* and *CNBP* genes, respectively, sequester MBNL proteins and thereby reduce their functional availability^47^. As a consequence, numerous clinical manifestations arise from disruption of the MBNL-regulated RNA processing program in tissues critically dependent on MBNL activity, including the eye, skeletal muscle, heart, nervous system, and immune cells^48^.

While the downstream consequences of MBNL dysfunction have been extensively studied, little is known about the mechanisms regulating expression of the predominant family member, *MBNL1*. To date, *MBNL1* regulation has largely been attributed to miRNA-dependent mechanisms and an autoregulatory feedback pathway^49, 50^. Elucidating the mechanisms that determine MBNL1 abundance is likely to be critical for understanding both physiological RNA regulatory networks during development or stress responses and pathological states associated with aberrant *MBNL1* expression, including cancer and repeat expansion disorders.

Here, we identify a previously unappreciated IPA pathway within the exceptionally long (∼115 kb) intron 1 of *MBNL1* that acts as a regulatory checkpoint controlling full-length transcript production in a tissue-and context-dependent manner. Through integrative transcriptomic analyses and experimental validation, we show that this pathway generates a stable polyadenylated isoform, hereafter called *MBNL1-IPA*, which is enriched in immune cells but strongly repressed in most other tissues and commonly used cell lines. We further demonstrate that splicing of the IPA-associated alternative last exon (ALE) is controlled by multiple regulatory layers. Reduced RNA polymerase II elongation modestly promotes ALE inclusion at the expense of canonical *MBNL1* expression, whereas depletion of *SFPǪ* strongly redirects transcript processing toward the *MBNL1-IPA* isoform in a manner dependent on the intron 1 context. Using antisense oligonucleotide (ASO) screening and minigene-based validation experiments, we identified a potent cis-regulatory element that functions as a ubiquitous molecular switch between the canonical and *MBNL1-IPA* isoform. Finally, we show that the *MBNL1* IPA pathway is activated by poly(I:C)-induced innate immune stress and contributes to stress-dependent reduction of MBNL1 abundance in both common cell lines and DM1-derived primary fibroblasts. Consistent with a direct role in this process, CRISPR-Cas9-mediated deletion of ALE counteracts stress-induced MBNL1 downregulation. Together, these findings uncover a previously unappreciated mechanism controlling MBNL1 dosage and establish the IPA pathway as a regulated checkpoint of *MBNL1* gene expression with implications for DM1 and other disorders associated with MBNL1 dysfunction.

## RESULTS

### Evidence for an active immune cell-enriched IPA pathway in the *MBNL1* locus

Given the regulatory potential of long introns, we investigated the ∼115 kb intron 1 of *MBNL1* for the presence of an intronic polyadenylation site (PAS). By integrating human poly(A)-selected RNA-seq datasets from multiple tissue types generated by ENCODE with PAS annotations from PolyA_DB, we identified a previously unappreciated intronic PAS located approximately 2.5 kb downstream of the first coding exon (e1) (**Figure 1A, Suppl. Figure S1A**). This PAS defines the 3′ end of a potential alternative last exon (ALE), which is predominantly present in immune cells and secondary lymphoid tissues (**Figure 1A**). To assess sequence conservation, we compared the ALE sequences across species and found its 3’ splice site (3’ss) to be only partially preserved in the murine *Mbnl1* locus, while no analogous human intronic PAS could be identified in mouse (**Suppl. Figure S1B-C**). Analysis of the *MBNL2* locus revealed a related intronic PAS (**Suppl. Figure S1D**), indicating that alternative 3′-end processing may represent a common regulatory feature of the human MBNL family.

**Figure 1.**
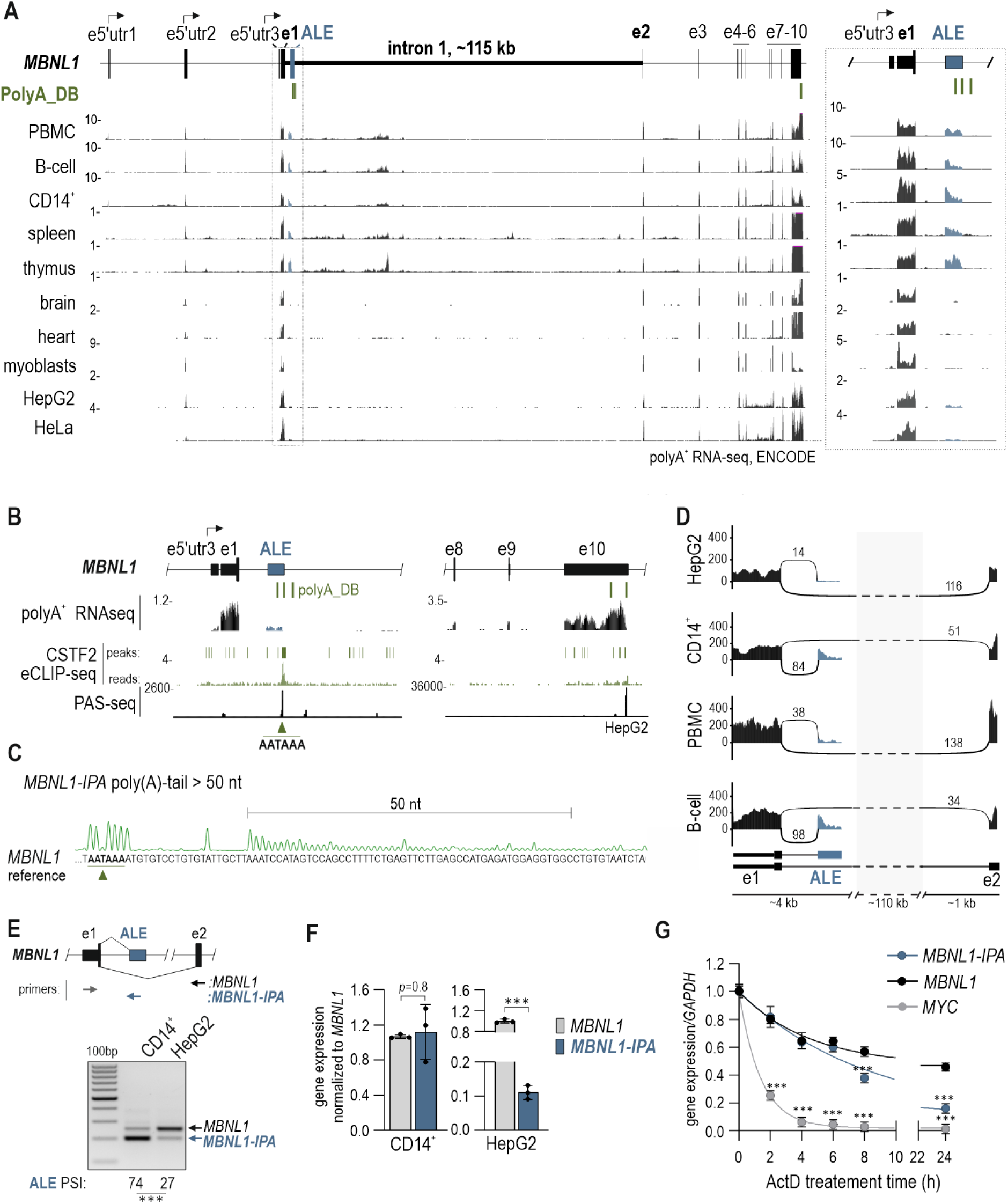
Evidence for an active immune cell-enriched IPA pathway in the MBNL1 locus. A) Schematic representation of RNA-processing events within the *MBNL1* locus inferred from the analysis of poly(A)+ RNA-seq datasets from ENCODE spanning multiple human tissues and cell types, together with annotated polyadenylation sites from PolyA_DB (green). Representative datasets are shown. Exons are drawn to scale, and arrows indicate alternative transcription start sites. The intronic last exon (ALE) associated with the annotated intronic PAS is highlighted in blue. The e1-ALE region (dashed box) is expanded on the right. B) Characterization of the intronic PAS within *MBNL1* intron 1. PAS-seq data from HepG2 cells (GSE218977)^51^, together with ENCODE CSTF2 eCLIP-seq tracks and PolyA_DB annotations, are displayed across the ALE region and the *MBNL1* 3’UTR. The putative ALE polyadenylation signal (green arrow) coincides with PAS-seq signal, PolyA_DB annotation, and a CSTF2 binding peak. C) Poly(A)-tail length analysis of *MBNL1-IPA* in THP1 cells. A representative Sanger sequencing trace of adenosines obtained from the poly(A)-tail assay is shown. The ALE polyadenylation signal (AATAAA) identified by poly(A)-tail mapping is indicated by the green arrow. The corresponding *MBNL1* genomic sequence is shown below. D) Sashimi plot analysis of canonical (e1-e2) and alternative (e1-ALE) splicing in CD14^+^ monocytes, B cells, PBMCs, and HepG2 cells using ENCODE RNA-seq datasets. Numbers indicate split-read counts supporting individual splice junctions. Canonical and ALE-associated splice junctions are shown in black and blue, respectively. E) Validation of canonical *MBNL1* and *MBNL1-IPA* transcripts by three-primer RT–PCR. *Top*, schematic representation of the analyzed *MBNL1* region with primer positions. *Bottom*, representative RT–PCR products obtained using primers specific for canonical (e1-e2) or alternative (e1-ALE) isoforms at a 1:1:0.5 (e1_F:ALE_R:e2_R) primers ratio. The *MBNL1-IPA* isoform is predominantly detected in CD14⁺ monocytes. *N =* 3. F) Relative abundance of *MBNL1-IPA* and canonical *MBNL1* isoforms in immune (CD14⁺ monocytes) and non-immune (HepG2) cells measured by RT–qPCR. *N =* 3. G) Stability of *MBNL1-IPA* assessed using actinomycin D in HEK293 cells. Transcript level was quantified by RT-qPCR at the indicated time points following transcriptional inhibition, relative to *GAPDH* and normalized to its level at T0. *MYC* served as a rapidly degraded control transcript. *N =* 3.

We next asked whether this candidate intronic PAS is actively recognized by the 3′-end processing machinery and gives rise to a distinct RNA species. First, we examined publicly available poly(A)-site sequencing (PAS-seq) samples from the immortalized human liver cell line (HepG2)^51^. We found substantial PAS-seq signal at the intronic PAS, reaching approximately 10% of that detected at the predominant *MBNL1* polyadenylation site within the canonical 3′UTR (**Figure 1B**). Because CSTF2 is recruited to functional PASs during 3′-end processing, we cross-referenced the intronic PAS coordinates with ENCODE enhanced crosslinking and immunoprecipitation (eCLIP-seq) datasets. We found a prominent CSTF2 binding site overlapping the intronic PAS and ALE boundaries, further supporting functional processing at this site (**Figure 1B**). We next sought to experimentally validate the resulting transcript, *MBNL1-IPA*, by analyzing its poly(A) tail and cleavage site. These analyses demonstrated that *MBNL1-IPA* is cleaved at a position consistent with PolyA_DB annotations, 23 nt downstream of the intronic PAS and carries a poly(A) tail exceeding 50 adenosines (**Figure 1C, Suppl. Figure S1E**).

To determine whether ALE acts solely as a terminal exon, we analyzed splice junction usage across intron 1 in RNA-seq datasets from immune cells and HepG2 cells. Our analysis revealed two mutually exclusive splicing outcomes: exon 1 joined either ALE or exon 2 (**Figure 1D, Suppl. Figure S1F**). Importantly, we found no evidence that ALE was subsequently spliced to downstream exons, supporting its designation as a terminal exon. We next asked how frequently the IPA pathway contributes to *MBNL1* transcript production across different cellular contexts. We estimated the relative representation of ALE-containing transcripts by calculating the proportion of exon 1 splice junctions connecting to ALE relative to exon 2. Immune cells exhibited substantially higher ALE inclusion (21-74%) than HepG2 cells (10%), indicating preferential expression of the alternative isoform in the immune lineage. To validate these findings experimentally, we performed RT-PCR using junction-specific primers that distinguish the canonical e1-e2 transcripts from the alternative e1-ALE isoform. While the canonical transcripts were readily detected in both HepG2 cells and monocytes (CD14^+^), the e1-ALE isoform was markedly enriched in monocytes (**Figure 1E**), consistent with the RNA-seq results. To quantitatively assess the proportion of *MBNL1-IPA* to canonical *MBNL1* isoforms, we analyzed their relative steady-state expression across a diverse panel of cell types. Consistent with transcriptomic analyses, *MBNL1-IPA* was present at nearly equivalent level to canonical *MBNL1* in monocytes, whereas it was 10-to 32-fold lower in commonly used cell lines and cancer derived immune cells (**Figure 1F, Suppl. Figure S1G**).

The defined cleavage site, long poly(A) tail and substantial representation of *MBNL1-IPA* in immune cells suggested that this transcript is unlikely to represent a transient by-product of premature transcription termination. We therefore examined the stability of *MBNL1-IPA* following treatment with the transcription inhibitor actinomycin *D (ActD)*. *MBNL1-IPA* was markedly more stable than the short-lived *MYC* transcript and showed modestly lower stability compared with canonical *MBNL1* mRNA (**Figure 1G**). Because *MBNL1-IPA* accumulates as a stable and abundant polyadenylated RNA, we next asked whether it might also engage the translational machinery. Polysome fractionation, ribosome profiling and epitope-tagging experiments collectively supported translation of a predicted micropeptide from *MBNL1-IPA* RNA (**Suppl. Figure S2**). However, the endogenous expression and biological relevance of the micropeptide remain unknown and were not explored by us further in the present study.

Collectively, these results establish the very long intron 1 as an important regulatory hub that expands *MBNL1* transcript diversity through cell type-specific alternative 3′-end processing.

### *MBNL1-IPA* is affected by MBNL-dependent exon 1 splicing

The detection of *MBNL1-IPA* isoforms originating from all three annotated *MBNL1* promoters suggests that *MBNL1-IPA* expression is primarily determined by downstream transcription and co-transcriptional processing rather than by promoter selection (**Suppl. Figure S3A,B**). We therefore asked whether *MBNL1-IPA* is subject to the well-established and MBNL-regulated alternative splicing of exon 1. To selectively deplete canonical *MBNL1* while preserving *MBNL1-IPA*, we used siRNA targeting the 3′UTR of the canonical isoform only (**Figure 2A, Suppl. Figure S3C**). Upon its depletion in HeLa cells, *MBNL1-IPA* abundance increased by 2-fold (**Figure 2A**). Conversely, MBNL1 overexpression reduced the abundance of both canonical exon 1-containing transcripts and *MBNL1-IPA* (**Figure 2B, Suppl. Figure S3D**).

**Figure 2.**
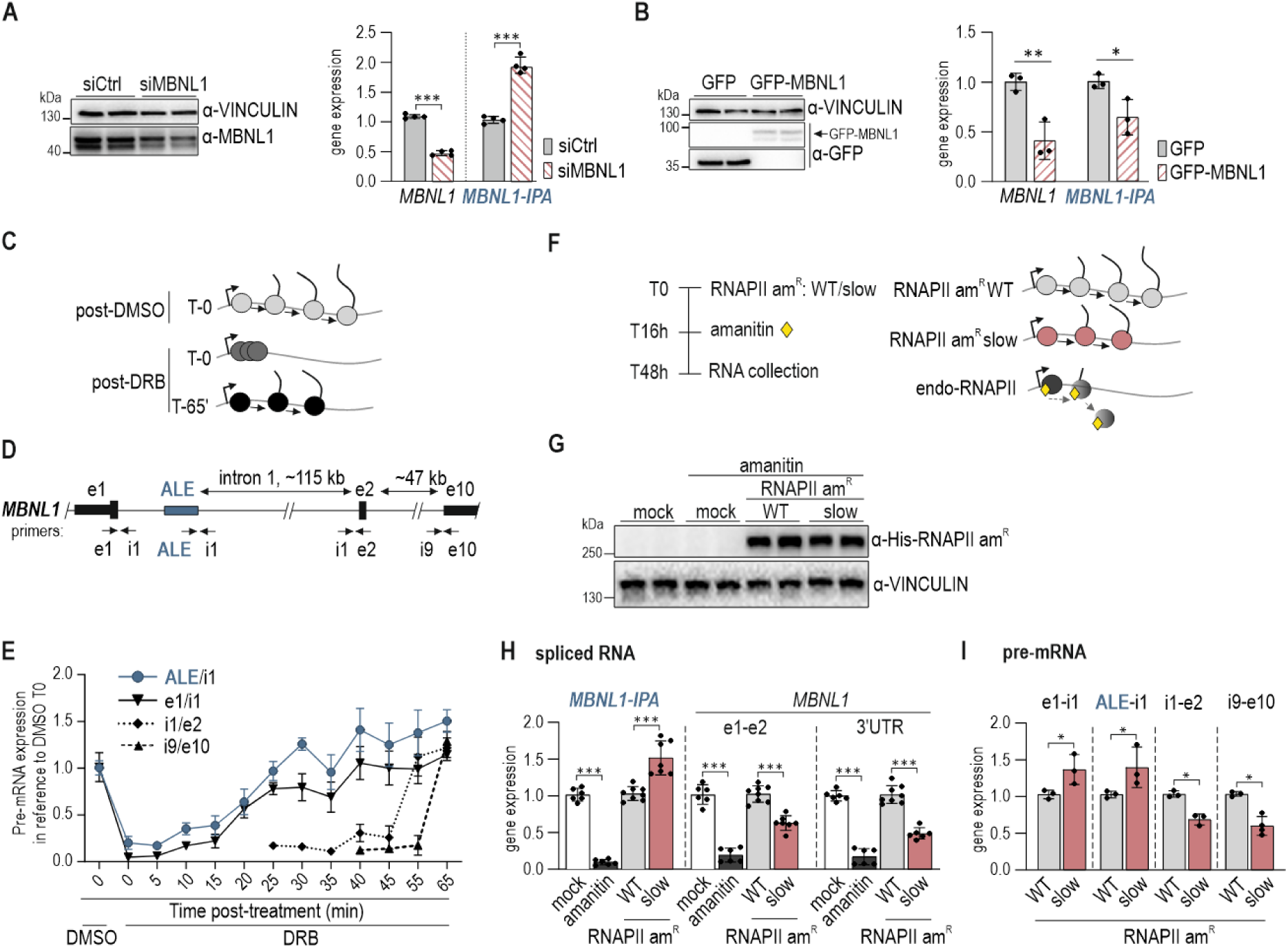
MBNL-dependent exon 1 splicing and transcription kinetics contribute to *MBNL1-IPA* expression. A) Effect of canonical *MBNL1* depletion on *MBNL1-IPA* expression in HeLa cells. *Left*, representative western blot showing MBNL1 depletion upon siMBNL1 treatment. VINCULIN served as a loading control. *Right*, RT–qPCR analysis of canonical *MBNL1* and *MBNL1-IPA* isoforms relative to *GAPDH* and normalized to siCtrl-treated cells. *N* = 3. B) Effect of MBNL1 overexpression on an *MBNL1-IPA* level in HeLa cells. *Left*, representative western blot showing GFP-MBNL1 or control GFP overexpression in HeLa cells. VINCULIN served as a loading control. *Right*, expression level of endogenous *MBNL1-IPA* and *MBNL1* isoforms containing exon 1 assessed by RT-qPCR analyses using a forward primer specific for 5’UTR within e1 which is absent in GFP-MBNL1 expressing vector. Data are presented relative to *TBP* and normalized to GFP-transfected cells. *N =* 3 C) Schematic of the DRB release assay used to monitor transcriptional progression across the *MBNL1* locus. Cells were treated with DRB to transiently inhibit RNA polymerase II (RNAPII) elongation and collected at the indicated time points following DRB washout. DMSO-treated cells served as a control. D) Schematic representation of the *MBNL1* locus showing the positions of RT-qPCR primers used to monitor RNAPII progression across intron 1 and downstream regions. Exons are shown as boxes and introns as lines, and primers are shown as arrows. Distances between selected exons are indicated. E) Analysis of *MBNL1* transcriptional progression following DRB release. Nascent pre-mRNA was quantified by RT-qPCR using primers spanning e1-i1, ALE-i1, i1-e2, and i9-e10. Transcript levels are expressed relative to DMSO-treated cells at T0. *N =* 3. F) Experimental design for modulation of RNAPII elongation kinetics. Cells were transfected with α-amanitin-resistant (am^R^) wild-type (WT) or slow RNAPII variants followed by inhibition of endogenous RNAPII using α-amanitin. Cells were analyzed 48 h after transfection. G) Expression of α-amanitin-resistant RNAPII am^R^ variants. Overexpressed WT and slow RNAPII am^R^ proteins were detected by western blot using anti-His-tag antibodies. VINCULIN served as a loading control. H) RT-qPCR analysis of *MBNL1-IPA* and canonical *MBNL1* transcripts in cells expressing WT or slow RNAPII am^R^. Canonical *MBNL1* was quantified using e1-e2 and 3′UTR amplicons. Expression levels are shown relative to *GAPDH* and normalized to WT RNAPII am^R^ -expressing cells. *N =* 6-8. I) Distribution of *MBNL1* pre-mRNA across the locus in cells expressing WT or slow RNAPII am^R^. Transcript abundance was measured by RT-qPCR using e1-i1, ALE-i1, i1-e2, and i9-e10 amplicons relative to *GAPDH* and normalized to WT RNAPII am^R^ -expressing cells. *N =* 3.

Together, these results demonstrate that *MBNL1-IPA* is subject to MBNL-dependent exon 1 autoregulation and responds dynamically to changes in MBNL1 abundance.

### RNA polymerase II elongation kinetics regulates IPA-driven exonization

We next investigated how processing of *MBNL1* intron 1 contributes to ALE activation and *MBNL1-IPA* expression. We first examined the contribution of RNA polymerase II (RNAPII) transcriptional dynamics. RNAPII elongation kinetics is a well-established determinant of alternative RNA processing because it influences the temporal window available for recognition of splice sites and polyadenylation sites, including intronic PASs^52^. In particular, reduced elongation rates can favor utilization of weak splice sites and promote IPA pathways within long genes^14, 18, 53^.

As an initial step, we evaluated the strength of the ALE cryptic 3′ss using MaxEntScan^54^. Despite its intronic location, the ALE acceptor site scored 8, well within the range of functional mammalian 3’ss. However, its predicted strength was substantially lower than that of the canonical e2 acceptor site (score 11), suggesting that ALE recognition may be particularly sensitive to transcription kinetics (**Suppl. Figure S3E**).

To determine the time required for RNAPII to transcribe across intron 1, we performed a DRB (5,6-dichloro-1-β-D-ribofuranosylbenzimidazole) release assay in HEK293 cells. Transient inhibition of transcription elongation by DRB enabled synchronized RNAPII release upon drug removal, and its progression through the *MBNL1* locus which we monitored by RT-qPCR using primer pairs distributed across *MBNL1* pre-mRNA (**Figure 2C,D**). The region encompassing ALE was transcribed shortly after DRB release, whereas RNAPII required more than 45 min to reach e2 and downstream portions of the gene (**Figure 2E**). We reasoned that slower RNAPII elongation would extend the temporal window between transcription of e1 and ALE, thereby favoring utilization of the weaker ALE 3′ss. To test this directly, we depleted endogenous RNAPII using α-amanitin and expressed α-amanitin-resistant (am^R^) *POLR2A* variants encoding the largest catalytic subunit of RNAPII, with either normal (RNAPII am^R^ WT) or approximately 3-fold slower elongation kinetics (RNAPII am^R^ slow) (**Figure 2F,G ; Suppl. Figure S3F,G**).

Expression of the slow RNAPII am^R^ variant increased ALE inclusion while reducing the abundance of canonical *MBNL1* transcripts measured at both the e1-e2 junction and the 3′UTR (**Figure 2H**), indicating a shift toward IPA-mediated RNA processing.

To assess whether increased ALE inclusion alters transcriptional progression through the *MBNL1* locus, we measured pre-mRNA abundance at three different positions across intron 1 and near the final canonical exon. Cells expressing the slow RNAPII am^R^ variant displayed an elevated pre-mRNA level across the proximal portion of intron 1, including the ALE region, together with reduced signal at e2 and the 3’UTR (**Figure 2I**). This redistribution is consistent with delayed transcriptional progression through the *MBNL1* locus and, together with increased ALE inclusion, supports a model in which slower elongation favors engagement of the *MBNL1* IPA pathway at the expense of full-length *MBNL1* transcript production.

Overall, these findings identify RNAPII elongation kinetics as an important determinant of *MBNL1* IPA and suggest that efficient transcription through the long intron 1 is a key mechanism restraining the IPA pathway.

### SFPQ suppresses activation of the *MBNL1* IPA pathway

Given the sensitivity of the *MBNL1* IPA pathway to elongation kinetics, we next asked whether RNA-binding proteins contribute to maintaining repression of ALE within the intron 1 of *MBNL1*. Several RBPs, including hnRNPM^55^, hnRNPC^56^, PTBP1^56^ and SFPQ^15, 57^, have been implicated in the suppression of cryptic RNA-processing events and maintenance of integrity of long-gene pre-mRNAs. We therefore examined whether these factors influence RNA processing at the *MBNL1* locus.

As an initial screen, we analyzed publicly available ENCODE RNA-seq datasets generated after 6 days of shRNA-mediated depletion of these RBPs in HepG2 cells. Among the tested factors, *SFPǪ* depletion produced the strongest reduction in RNA-seq coverage downstream of exon 1 (**Figure 3A).** The substantial loss of downstream transcript signal, together with the established role of SFPQ in preserving long-gene transcription^57, 58^, prompted us to investigate this factor further.

**Figure 3.**
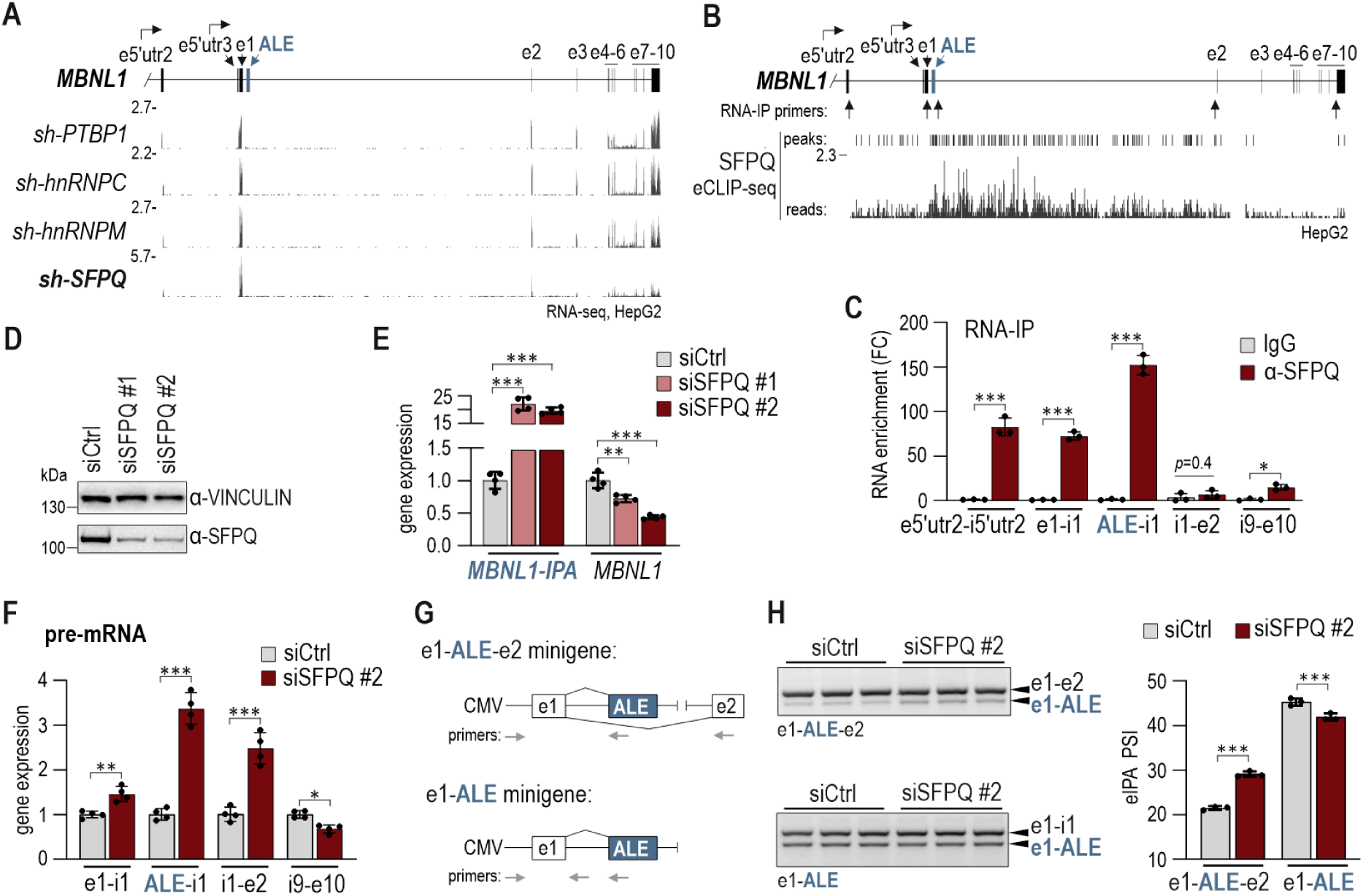
SFPQ suppresses activation of the *MBNL1* IPA pathway. A) RNA-seq coverage profiles across the *MBNL1* locus following shRNA-mediated depletion of *hnRNPM*, *hnRNPC*, *PTBP1*, or *SFPǪ* in HepG2 cells (ENCODE). Exons are drawn to scale and arrows indicate alternative transcription start sites. B) Schematic representation of the *MBNL1* locus showing SFPQ binding sites identified by ENCODE eCLIP-seq in HepG2 cells. Arrows indicate transcript regions analysed by RNA immunoprecipitation (RNA-IP) in panel C. C) RNA-IP analysis of SFPQ association with *MBNL1* transcript in HeLa cells. Immunoprecipitated RNA obtained using anti-SFPQ antibody (EPR11847) was quantified by RT-qPCR using amplicons indicated in panel B. Enrichment is shown relative to IgG control. *N =* 3. D) Validation of *SFPǪ* depletion in HeLa cells using two independent siRNA approaches: siSFPQ #1 (single siRNA) and siSFPQ #2 (siRNA pool). SFPQ protein abundance was assessed by western blot. VINCULIN served as a loading control. E) Effect of *SFPǪ* depletion on *MBNL1* transcript output. *MBNL1-IPA* and canonical *MBNL1* transcript levels were quantified by RT-qPCR and normalized to siCtrl-treated cells. *N =* 4. F) Distribution of *MBNL1* pre-mRNA across the locus following *SFPǪ* depletion. Transcript abundance was measured by RT-qPCR using amplicons spanning e1–i1, ALE–i1, i1–e2 and i9–e10 and normalized to siCtrl-treated cells. *N =* 4. G) Schematic representation of the e1–ALE–e2 and e1–ALE minigene reporters. Canonical exons, ALE, and position of primers used for transcript analyses are indicated. H) *Left*, RT–PCR analysis of transcripts generated from the e1–ALE–e2 and e1–ALE reporters in control and SFPQ-depleted HeLa cells. Canonical and ALE-containing PCR products are indicated. *Right*, quantification of ALE-containing isoforms, presented as percent spliced in (PSI). *N =* 4.

We first examined whether SFPQ associates with the *MBNL1* transcript. Analysis of ENCODE eCLIP datasets from HepG2 cells revealed extensive SFPQ binding across intron 1, with the strongest enrichment observed near its proximal region (**Figure 3B**). Consistent with these data, RNA immunoprecipitation performed in HeLa cells using two independent SFPQ antibodies reproducibly enriched the proximal region of the *MBNL1* transcript encompassing e5′utr2, e1 and ALE, whereas little enrichment was detected at e2 or e10 (**Figure 3B,C, Suppl. Figure S4A**). These findings indicate preferential association of SFPQ with the proximal region of the *MBNL1* pre-mRNA.

To determine the functional consequences of short-term but efficient SFPQ depletion, HeLa cells were transfected with either an individual siRNA or a pooled siRNA set targeting *SFPǪ*, both of which efficiently reduced *SFPǪ* expression at the RNA and protein levels (**Figure 3D, Suppl. Figure S4B,C**). *SFPǪ* silencing resulted in robust activation of the IPA pathway, leading to an approximately 20-fold increase in *MBNL1-IPA* level accompanied by a modest reduction in the canonical *MBNL1* mRNA (**Figure 3E**). Notably, even subtle reduction of *SFPǪ* was sufficient to significantly increase an *MBNL1-IPA* level, by 3-fold, without affecting canonical *MBNL1* mRNA (**Suppl. Figure S4D-F**), indicating that the IPA pathway is very sensitive to SFPQ dosage.

To assess the effect of SFPQ on transcriptional progression along the *MBNL1* locus, we quantified pre-mRNA abundance across intron 1 and downstream regions following silencing of *SFPǪ*. Loss of SFPQ resulted in accumulation of intron 1-containing pre-mRNA, accompanied by a slight reduction in distal transcript signal (**Figure 3F, Suppl. Figure S4G**), consistent with impaired productive transcription through the locus and enhanced engagement of the IPA pathway. Conversely, overexpression of SFPQ further suppressed *MBNL1-IPA* expression (**Suppl. Figure S4H-I**), supporting a direct role for SFPQ in regulating RNA processing across *MBNL1* intron 1 and repressing the IPA pathway.

Because SFPQ functions as a splicing regulator, it could potentially influence recognition of the ALE 3′ss. To test this possibility, we generated two minigene reporters: a longer reporter containing exon 1, the ALE region, exon 2 and its upstream intron, 1000 bp long (e1-ALE-e2), as well as a shorter reporter terminating shortly downstream of ALE (e1-ALE) (**Figure 3G**). Relative to control conditions, *SFPǪ* knockdown significantly enhanced ALE inclusion in the longer reporter, as evidenced by an increased percent spliced in (PSI) value, whereas the shorter construct predominantly accumulated the unspliced transcript (**Figure 3H**). Notably, both reporters contain the identical ALE 3′ss, yet enhanced ALE inclusion upon *SFPǪ* depletion was observed only in the long reporter. These findings suggest that SFPQ does not primarily act through local regulation of the ALE 3′ss but instead represses the IPA pathway through mechanisms that depend on the broader intronic context.

Taken together, these findings identify SFPQ as a key regulator of RNA processing across *MBNL1* intron 1 and maintenance of canonical *MBNL1* expression. Our data support a model in which SFPQ strongly suppresses the *MBNL1* IPA pathway through mechanisms that depend on the long-intron context.

### An intronic cis-regulatory element and the ALE region shape *MBNL1* expression through the IPA pathway

Having established that transcriptional dynamics and long-intron processing contribute to suppression of ALE inclusion, we next asked whether *MBNL1* intron 1 contains local cis-acting elements that regulate the IPA pathway.

To identify such elements, we performed a screening of antisense oligonucleotide blockers (ASO) targeting sequences surrounding the ALE-associated 3′ss. Nine overlapping ASOs spanning approximately 100 nucleotides were designed and evaluated using a three-primer RT-PCR assay (**Figure 4A**). Among the tested oligonucleotides, ASO-101 markedly increased ALE inclusion (**Figure 4B**). This effect was subsequently confirmed in four distinct cell lines where ASO-101 robustly increased *MBNL1-IPA* expression by 10-30-fold while substantially reducing the canonical *MBNL1* expression (**Figure 4C, Suppl. Figure S5A**). The magnitude, reproducibility and cell type-independence of this response pointed to the ASO-101-responsive region as a candidate cis-regulatory element controlling engagement of the IPA pathway.

**Figure 4.**
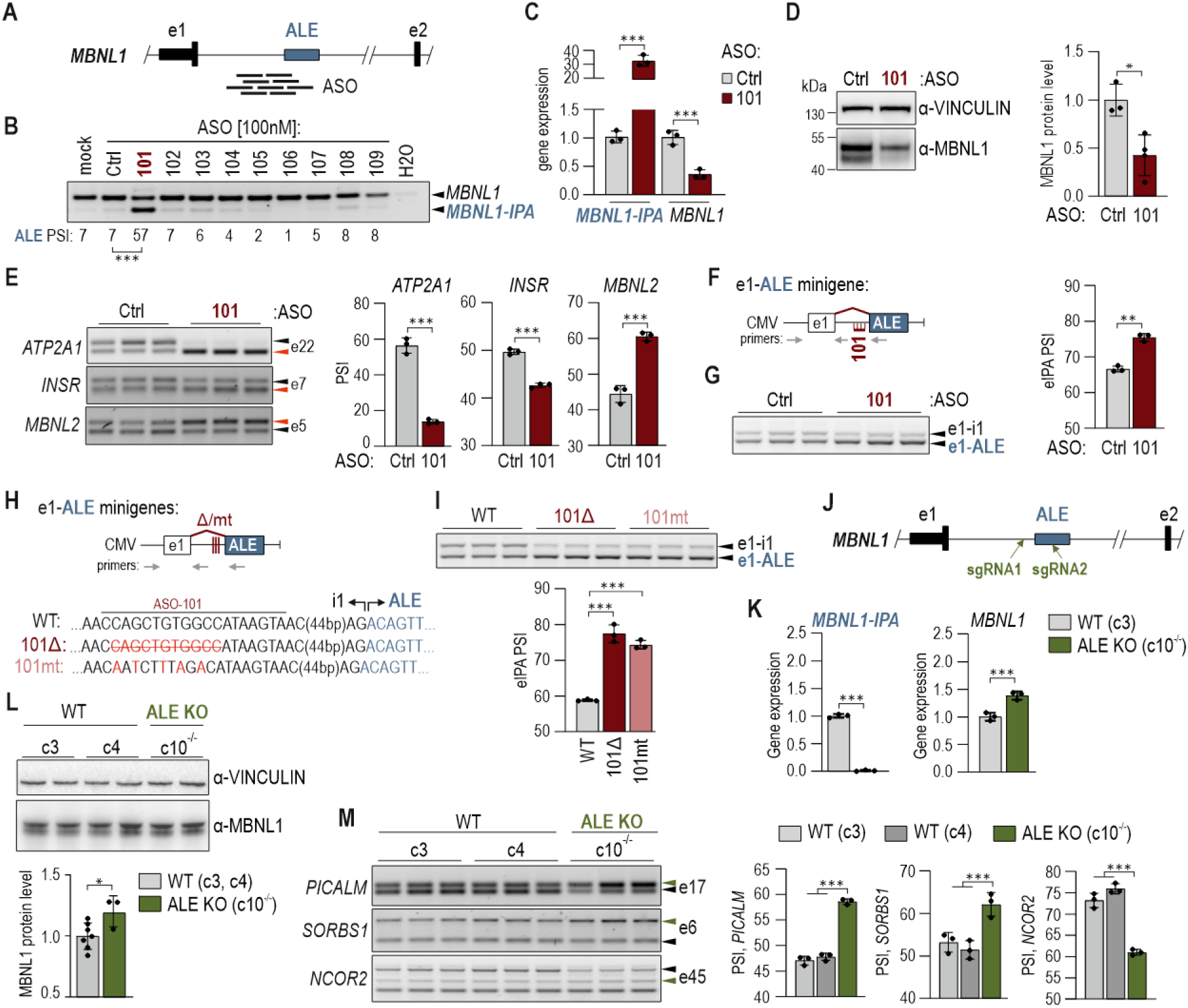
An intronic cis-regulatory element and the ALE region shape *MBNL1* expression through the IPA pathway. A) Schematic representation of the *MBNL1* exon 1–ALE-e2 region showing the position of antisense oligonucleotides (ASOs) targeting sequences upstream of and within ALE. B) Identification of an ALE regulatory region by ASO screening. Three-primer RT-PCR analysis of *MBNL1* isoforms following transfection of HeLa cells with individual ASOs. Canonical *MBNL1* and *MBNL1-IPA* isoforms are indicated. Below, quantification of ALE inclusion presented as PSI values normalized to ASO-Ctrl-treated cells. Statistically significant activation of ALE was observed only following ASO-101 treatment. *N =* 2. C) Effect of ASO-101 on *MBNL1* transcript output. Canonical *MBNL1* and *MBNL1-IPA* transcript levels were quantified by RT-qPCR and normalized to ASO-Ctrl-treated HeLa cells. *N =* 3. D) Effect of ASO-101 on MBNL1 protein abundance. *Left*, representative western blot analysis of MBNL1 protein abundance in HeLa cells treated with ASO-Ctrl and ASO-101. VINCULIN served as a loading control. *Right*, quantification of MBNL1 protein abundance relative to VINCULIN and normalized to ASO-Ctrl-treated cells. *N =* 3-4. E) Functional consequences of ASO-101 treatment on MBNL-dependent splicing. *Left*, RT–PCR analysis of *ATP2A1* exon 22, *INSR* exon 7, and *MBNL2* exon 5. Isoforms associated with reduced MBNL1 activity are indicated by red arrows. *Right*, quantification of splicing pattern of MBNL1-dependent events shown as PSI values. *N =* 3. F) Schematic representation of the e1–ALE minigene reporter. Canonical exons, ALE, the ASO-101 targeted region, and primer positions used for transcript analysis are indicated. G) Response of the e1-ALE reporter to ASO-101 treatment in HeLa cells. *Left*, RT–PCR analysis of transcripts generated from the reporter in ASO-Ctrl and ASO-101-treated cells. Unspliced (e1-i1) and e1-ALE spliced RNAs are indicated. *Right*, quantification of ALE inclusion presented as PSI values in reference to unspliced transcript. *N =* 3. H) Schematic representation of WT and mutant e1–ALE minigene reporters. The deleted GC-rich sequence in the 101Δ mutant variant is indicated in red and crossed out, while nucleotide substitutions introduced into the ASO-101-targeted region in the 101mt mutant variant are highlighted in red. Primer positions used for transcript analysis are shown. I) Effect of GC-rich element mutations in the e1-ALE reporter on ALE inclusion. *Left*, RT–PCR analysis of transcripts generated from WT, 101Δ, and 101mt reporters. Unspliced (e1-i1) and e1-ALE spliced RNAs are indicated. *Right*, quantification of ALE inclusion presented as PSI values in reference to unspliced transcript. *N =* 3. J) Schematic representation of the *MBNL1* exon 1–ALE-e2 fragment showing the approximate positions of sgRNAs used for CRISPR-Cas9-mediated deletion of part of the ALE region. K) RT–qPCR quantification of *MBNL1-IPA* and *MBNL1* isoforms in the WT HEK293 clone (c3) and ALE KO clone (c10^-/-^) which carries a homozygous deletion. Expression level is normalized to the WT clone c3. *N =* 3. L) *Left*, representative western blot analysis of MBNL1 protein abundance in two WT clones (c3 and c4) and ALE KO clones (c10^-/-^). VINCULIN served as a loading control. *Right*, quantification of MBNL1 protein abundance, relative to VINCULIN and normalized to WT clones (c3 and c4). Statistical significance was determined relative to two WT clones (c3 and c4). *N =* 3-4. M) Functional consequences of ALE deletion on MBNL-dependent splicing. *Left*, RT–PCR analysis of *PICALM* e17, *SORBS1* e6, and *NCOR2* e45. Isoforms promoted by MBNL1 are indicated by green arrows. *N =* 3. *Right*, quantification of alternative splicing events, presented as PSI values for the indicated alternative exons in control and ALE KO cell lines. Statistical significance was normalized to WT clones (c3 and c4). *N =* 3.

Consistent with the reduction in canonical isoforms, ASO-101 treatment led to a 2-fold decrease in MBNL1 protein abundance (**Figure 4D**), whereas expression of the closely related paralog MBNL2 remained unaffected (**Suppl. Figure S5B**). To assess downstream functional consequences of ASO-101-mediated activation of the IPA pathway, we analyzed three established MBNL-dependent splicing events, including the positively regulated *ATP2A1* e22 and *INSR* e7, and the negatively regulated *MBNL2* e5. In each case, ASO-101 induced splicing changes consistent with reduced MBNL1 activity, demonstrating that activation of the IPA pathway is sufficient to alter MBNL-dependent RNA processing (**Figure 4E**).

To determine whether the regulatory activity of the ASO-101-responsive region requires a long-intron context, we examined the previously established e1-ALE and e1-ALE-e2 minigene reporters (**Figure 4F, Suppl. Figure S5C**). Both reporters responded to ASO-101 treatment with increased ALE inclusion (**Figure 4G, Suppl. Figure S5D**), indicating that regulation by this element can be recapitulated outside the endogenous locus and does not strictly depend on the surrounding long-intron context.

To gain insight into its mechanism of action, we searched for putative RBP-binding motifs within the ASO-101-targeted region. Analysis using SpliceAid software^59^ did not reveal high-confidence binding sites for known splicing regulators (**Suppl. Figure S5E**). Instead, the region displayed a pronounced GC-rich sequence composition, raising the possibility that its regulatory activity depends on local sequence or RNA structural features rather than a direct RBP-mediated interaction. Although the latter cannot be ruled out. To test this hypothesis, we generated two minigene variants: one containing an 11-nt deletion within the GC-rich region (101Δ) and another carrying five A/T substitutions (101mt) (**Figure 4H, Suppl. Figure S5F**). Both perturbations recapitulated the effect of ASO-101, increasing ALE inclusion by approximately 20% (**Figure 4I, Suppl. Figure S5G).** The concordant effects of ASO masking, targeted deletion, and sequence substitution support a direct regulatory role for this GC-rich intronic element in repression of the IPA pathway.

We next asked whether genetic disruption of the IPA pathway could redirect RNA processing toward productive *MBNL1* expression and thereby enhance MBNL-dependent splicing, even in cells with minimal basal ALE inclusion. To address this possibility, we generated a CRISPR–Cas9-edited HEK293 cell line harboring a deletion encompassing the ALE-associated 3′ss and the newly identified cis-regulatory element (**Figure 4J, Suppl. Figure S6A-D**). We obtained a homozygous clone (ALE KO c10^-/-^) exhibiting complete loss of *MBNL1-IPA* expression but increased level of canonical *MBNL1* isoforms at both the RNA and protein levels (**Figure 4K, L**). To determine whether this increase translated into functional consequences, we examined well-characterized MBNL-regulated alternative splicing events in ALE KO and two control cell lines (clone 3 and 4). Nine of the twelve events displayed splicing changes in the expected MBNL-dependent direction, including seven MBNL-activated and two MBNL-repressed exons, while three remained unchanged (**Figure 4M, Suppl. Figure S6E,F).** Notably, these splicing changes occurred despite only a modest increase in MBNL1 protein abundance, indicating that suppression of the IPA pathway is sufficient to measurably enhance MBNL-dependent splicing.

Collectively, these findings identify a potent intronic cis-regulatory element that suppresses the *MBNL1* IPA pathway and serves as a major local determinant of *MBNL1* transcript output. Disruption of this element by antisense masking or mutagenesis redirects RNA processing toward the *MBNL1-IPA* isoform and reduces MBNL1 activity.

Moreover, our results demonstrate that the *MBNL1* IPA pathway exerts a measurable constitutive influence on endogenous *MBNL1* expression even in cells in which it is strongly suppressed under basal conditions.

### Poly(I:C)-induced immune stress remodels *MBNL1* transcript output through engagement of the IPA pathway

Given the established link between cellular stress and increased utilization of intronic PAS^60^, we investigated whether stress signaling engages the *MBNL1* IPA pathway.

To address this question, HeLa cells were subjected to endoplasmic reticulum stress (ER) induced by thapsigargin (TG), oxidative stress induced by hydrogen peroxide (H₂O₂) or sodium arsenite (NaAsO₂), and innate immune stress induced by poly(I:C), a synthetic double-stranded RNA that mimics viral infection (**Figure 5A**). *MBNL1-IPA* expression was quantified following acute stress exposure. Among the tested conditions, poly(I:C) elicited the strongest response, indicating that innate immune activation is a potent inducer of the *MBNL1* IPA pathway (**Figure 5B**).

Given the preferential expression of *MBNL1-IPA* in immune cells, we further characterized the response of *MBNL1* locus to 6 h exposure to poly(I:C). Efficient activation of innate immune signaling was confirmed by robust induction of Interleukin-8 expression, *CXCL8* (**Figure 5C**). Poly(I:C) treatment produced a modest increase in *MBNL1* transcriptional activity, measured at the 5′ region of the transcript (**Figure 5D**). To account for stress-induced activation of the *MBNL1* locus, steady-state levels of *MBNL1-IPA* and canonical *MBNL1* transcripts were referenced to an *MBNL1* pre-mRNA level. Relative to this measure of locus activity, poly(I:C) treatment resulted in an approximately 3-fold enrichment of *MBNL1-IPA* and a concomitant tendency toward reduced canonical *MBNL1* expression (**Figure 5E**). These findings indicate that innate immune stress activates the *MBNL1* locus while redirecting transcript output toward the *MBNL1-IPA* isoform in HeLa cells.

We next examined whether activation of the IPA pathway translates into altered MBNL1 protein abundance. Following 6 h poly(I:C) treatment and an additional 18 h post-stress induction (PtSI) period (**Figure 5F**), MBNL1 protein abundance decreased by approximately 25% (**Figure 5G**). MBNL2 protein level was also reduced and accompanied by alterations in *MBNL2* RNA processing (**Suppl. Figure S7A-D**), suggesting that innate immune activation affects MBNL homeostasis more broadly.

Because innate immune stress triggers extensive transcriptomic and proteomic remodeling^61^, we sought to determine the contribution of the *MBNL1* IPA pathway to stress-induced changes in MBNL1 protein abundance. To test this, WT and ALE KO HEK293 cells were subjected to the same poly(I:C) treatment protocol as used in HeLa cells. Whereas WT cells exhibited a slight reduction in MBNL1 protein abundance, ALE KO cells instead showed increased MBNL1 protein abundance following stress exposure (**Figure 5H**).

Together, these results are consistent with the *MBNL1* IPA pathway contributing to innate immune stress-dependent modulation of *MBNL1* expression.

### Poly(I:C)-induced innate immune stress exacerbates MBNL1 dysfunction in DM1-affected fibroblasts

Given that innate immune activation promotes the IPA pathway while reducing MBNL1 expression, we asked whether immune stress could further exacerbate the MBNL loss-of-function molecular phenotype in DM1. To address this question, we used fibroblasts derived from a DM1 patient carrying a heterozygous expansion of approximately 1,700 CTG repeats within the *DMPK* gene^62^. Primary fibroblasts are increasingly recognized as important contributors to innate immunity, directly sensing viral nucleic acids and activating antiviral and inflammatory signaling pathways^63, 64^.

Fibroblasts were exposed to poly(I:C) and analyzed either immediately after a 6 h treatment or following an additional 18 h PtSI period (**Figure 6A**). Poly(I:C) treatment triggered a strong innate immune response, as evidenced by ∼100-fold induction of *CXCL8* expression after 6 h (**Figure 6B**). Despite robust immune activation, *MBNL1* locus activity, measured by pre-mRNA abundance at the 5′ end of the gene, remained unchanged (**Figure 6C**). To relate mature transcript abundance to transcriptional activity of the *MBNL1* locus, canonical *MBNL1* and *MBNL1-IPA* levels were referenced to *MBNL1* pre-mRNA. While *MBNL1-IPA* expression was not significantly affected, a canonical *MBNL1* transcript level was reduced (**Figure 6D**), suggesting the involvement of post-transcriptional mechanisms influencing *MBNL1* transcript turnover.

**Figure 5.**
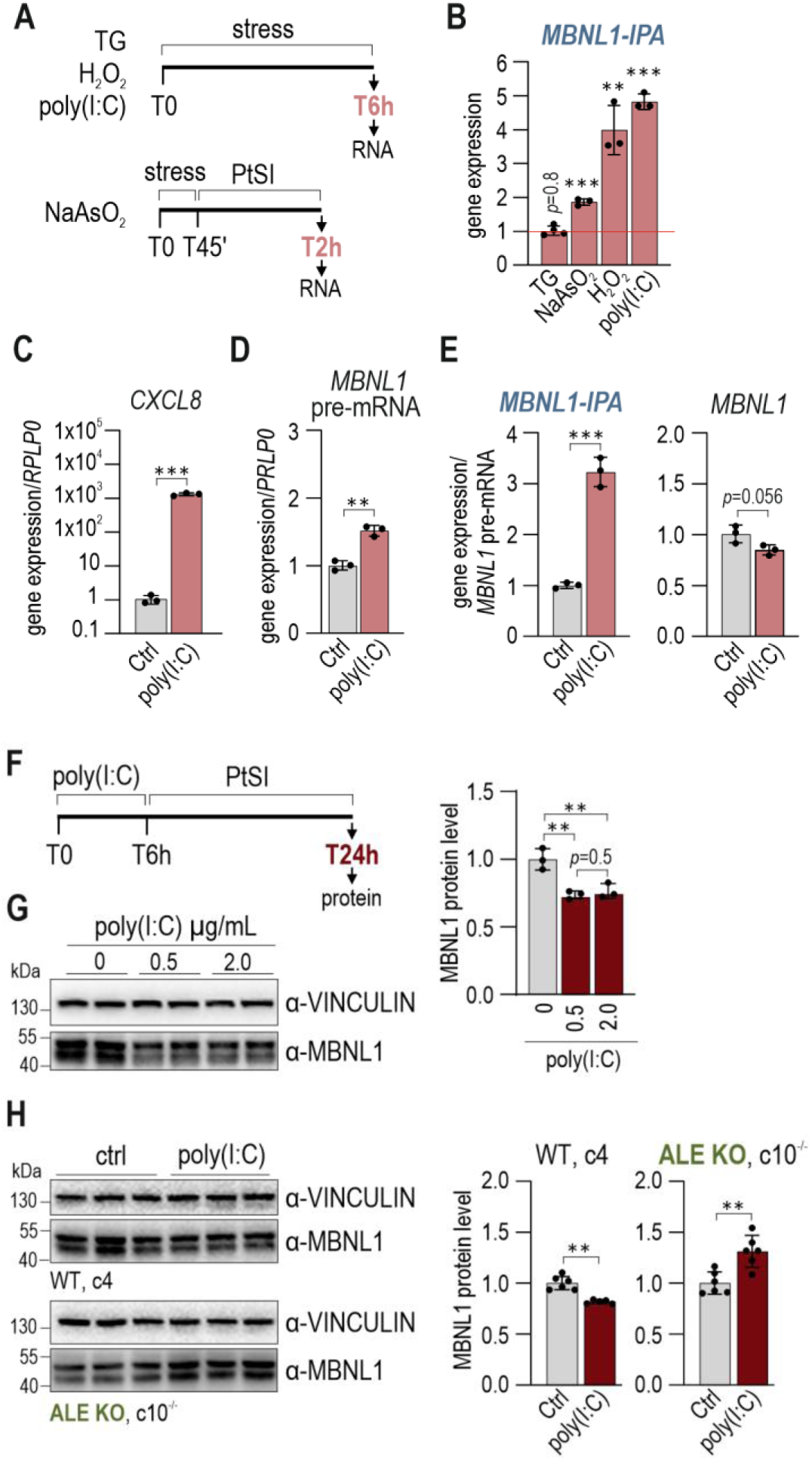
Poly(I:C)-induced immune stress remodels *MBNL1* transcript output through engagement of the IPA pathway. A) Experimental design of stress induction in HeLa cells. Cells were exposed to 3 µM thapsigargin (TG), 50 µM hydrogen peroxide (H₂O₂), 0.5 µg/mL poly(I:C), or 0.5 mM sodium arsenite (NaAsO₂) and collected at the indicated time points for RNA analysis. PtSI, post-stress induction. B) RT–qPCR quantification of *MBNL1-IPA* following the indicated stress induction. Expression levels are shown relative to various reference genes: *GAPDH* for TG and NaAsO_2_, *ACTB* for H_2_O_2_ and *RPLP0* for poly(I:C) and normalized to vehicle treated cells (H_2_O for TG, NaAsO_2_ and H_2_O_2_ or RNAiMAX-treated cells for poly(I:C)). *N =* 3-4. C) Validation of innate immune activation in HeLa cells following poly(I:C) treatment. *CXCL8* expression was quantified by RT–qPCR and normalized to RNAiMAX treated control cells. *N =* 3. D) Transcriptional activity of the *MBNL1* locus following poly(I:C) treatment. *MBNL1* pre-mRNA was quantified at the 5′ region of transcripts initiating from e5′UTR2 and normalized to RNAiMAX-treated control cells. *N =* 3. E) Redistribution of *MBNL1* transcript output following innate immune activation. Canonical *MBNL1* and *MBNL1-IPA* transcript levels were calculated in reference to *MBNL1* pre-mRNA shown in panel D and normalized to RNAiMAX-treated control cells. *N =* 3. F) Experimental design of the post-stress induction (PtSI) assay. HeLa cells were treated with poly(I:C) for 6 h and cultured in fresh medium for additional 18 h prior to protein analysis. G) Effect of poly(I:C)-induced innate immune stress on MBNL1 protein abundance in HeLa cells. *Left*, representative western blot. VINCULIN served as a loading control. *Right*, quantification of MBNL1 protein abundance relative to VINCULIN and normalized to RNAiMAX-treated control cells. *N =* 3. H) Contribution of the ALE element to stress-induced MBNL1 downregulation*. Left*, representative western blot analysis of MBNL1 abundance in WT (c4) and ALE KO (c10^-/-^) HEK293 cells following poly(I:C) treatment and PtSI. VINCULIN served as a loading control. *Right*, quantification of MBNL1 protein level in reference to VINCULIN and normalized to RNAiMAX treated control cells. *N =* 5-6.

In contrast, following the 18 h of PtSI, the inflammatory response became markedly amplified, with *CXCL8* expression reaching approximately 10^4^-fold induction (**Figure 6E**). This was accompanied by ∼3-fold increase in *MBNL1* transcriptional activity (**Figure 6F**). Relative to *MBNL1* pre-mRNA levels, transcript output was redistributed toward the IPA pathway, as evidenced by modest enrichment of *MBNL1-IPA* and reduced level of canonical *MBNL1* (**Figure 6G**). These findings indicate that prolonged innate immune signaling alters RNA-processing decisions at the *MBNL1* locus in DM1 fibroblasts.

**Figure 6.**
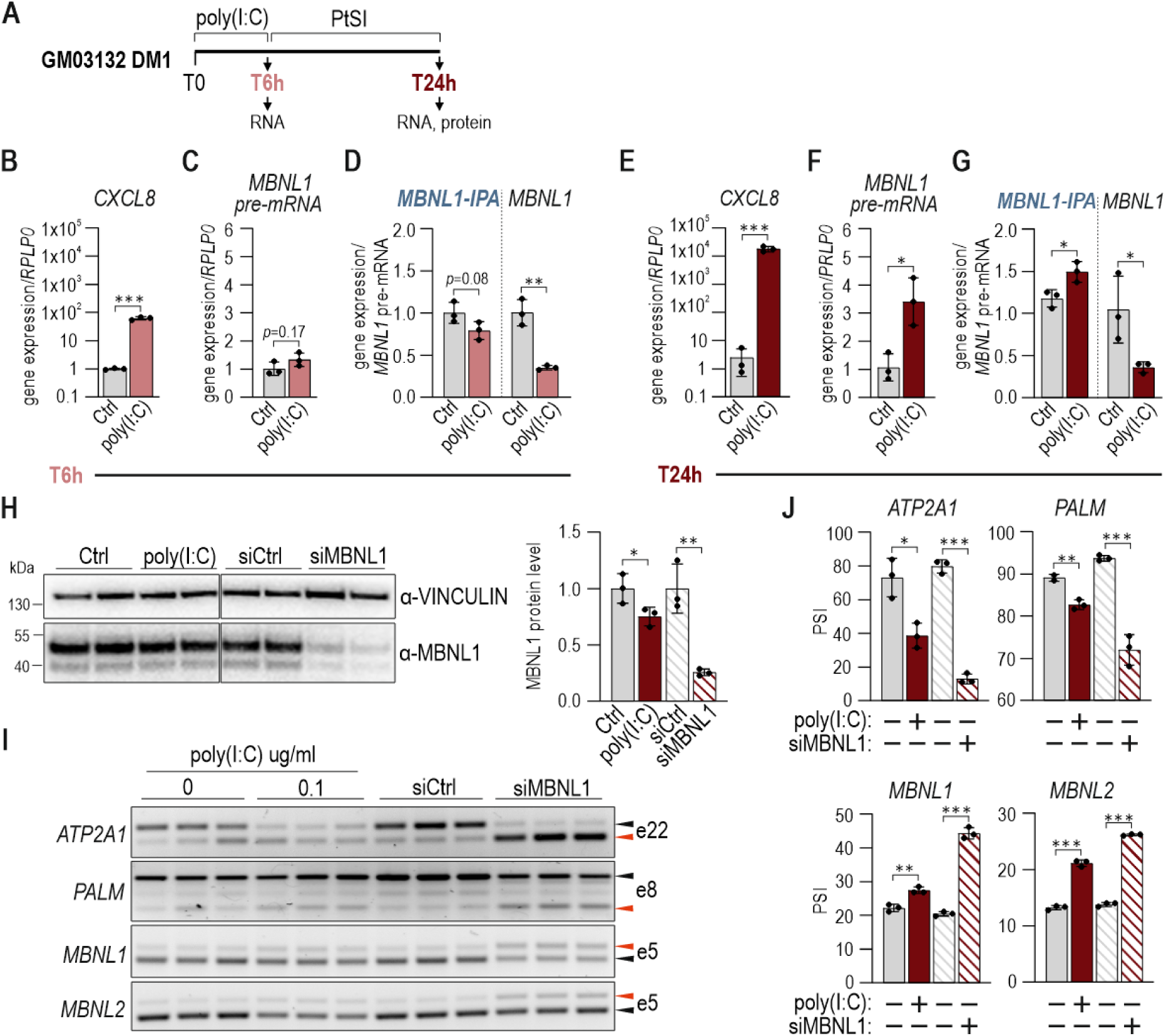
Poly(I:C)-induced innate immune stress reduces MBNL1 abundance and impairs MBNL-dependent splicing in DM1-affected fibroblasts. A) Experimental design of poly(I:C) treatment in GM03132 DM1 patient-derived fibroblasts. Cells were collected either following 6 h of poly(I:C) treatment or after additional 18 h of post-stress induction (PtSI) period. B) Validation of innate immune activation in DM1 fibroblasts. *CXCL8* expression was quantified by RT-qPCR following 6 h poly(I:C) treatment and normalized to RNAiMAX-treated control cells. *N =* 3. C) Transcriptional activity of the *MBNL1* locus following 6 h poly(I:C) treatment. *MBNL1* pre-mRNA was quantified at the 5′ region of transcripts initiating from e5′utr2 and normalized to RNAiMAX-treated control cells. *N =* 3. D) Effect of acute innate immune activation on *MBNL1* transcript output. Canonical *MBNL1* and *MBNL1-IPA* transcript levels were calculated in reference to *MBNL1* pre-mRNA shown in panel C and normalized to RNAiMAX-treated control cells. *N =* 3. E) As in panel B, but following poly(I:C) treatment and the 18 h PtSI period. F) As is panel C, but following poly(I:C) treatment and the 18 h PtSI period. G) As is panel D, but following poly(I:C) treatment and the 18 h PtSI period. H) Effect of poly(I:C)-induced innate immune stress on MBNL1 protein abundance in DM1-derived fibroblasts. *Left*, representative western blot analysis of MBNL1 protein abundance following poly(I:C) treatment and PtSI. VINCULIN served as a loading control. siMBNL1-treated cells were included as a positive control for *MBNL1* depletion. *Right*, quantification of MBNL1 protein abundance relative to VINCULIN and normalized to control conditions, either RNAiMAX-treated control cells or siCtrl-treated cells. *N =* 3. I) RT–PCR analysis of selected MBNL1-dependent splicing events *ATP2A1* e22, *MBNL1* e5, *MBNL2* e5 and *PALM* e8 in DM1-affected fibroblasts following poly(I:C) treatment and the 18 h PtSI period or siRNA-mediated knockdown of *MBNL1*. J) Quantification of alternative splicing events shown in panel I presented as PSI values. *N =* 3.

Consistent with the transcript-level changes, MBNL1 protein abundance decreased by approximately 20% following 6 h poly(I:C) treatment and 18 h PtSI period (**Figure 6H**). To assess the functional significance of this reduction, we compared the effects of stress exposure with those induced by siRNA-mediated *MBNL1* depletion (**Figure 6H**). Analysis of six established MBNL-dependent splicing events revealed that four exhibited splicing changes in the same direction as those observed following *MBNL1* knockdown, whereas the remaining two events were unaffected (**Figure 6I,J, Suppl. Figure S8A**).

Together, these results indicate that innate immune stress alters *MBNL1* transcript processing in DM1-affected fibroblasts, reduces MBNL1 protein abundance, and promotes a partial shift toward the MBNL loss-of-function splicing pattern.

## DISCUSSION

Our study identifies the IPA pathway as a novel mechanism controlling *MBNL1* expression in a context-dependent manner and in response to cellular stress. We show that IPA-associated splicing of ALE within the *MBNL1* intron 1 generates a short, relatively stable and polyadenylated isoform, *MBNL1-IPA*, and attenuates production of canonical *MBNL1* isoforms. Mechanistically, this pathway is regulated by regulatory factors, including RNAPII elongation kinetics, SFPQ and a potent local cis-regulatory element. Furthermore, we demonstrate that innate immune stress activates the IPA pathway, reduces MBNL1 abundance, and alters MBNL-dependent RNA processing (**Figure 7**).

**Figure 7.**
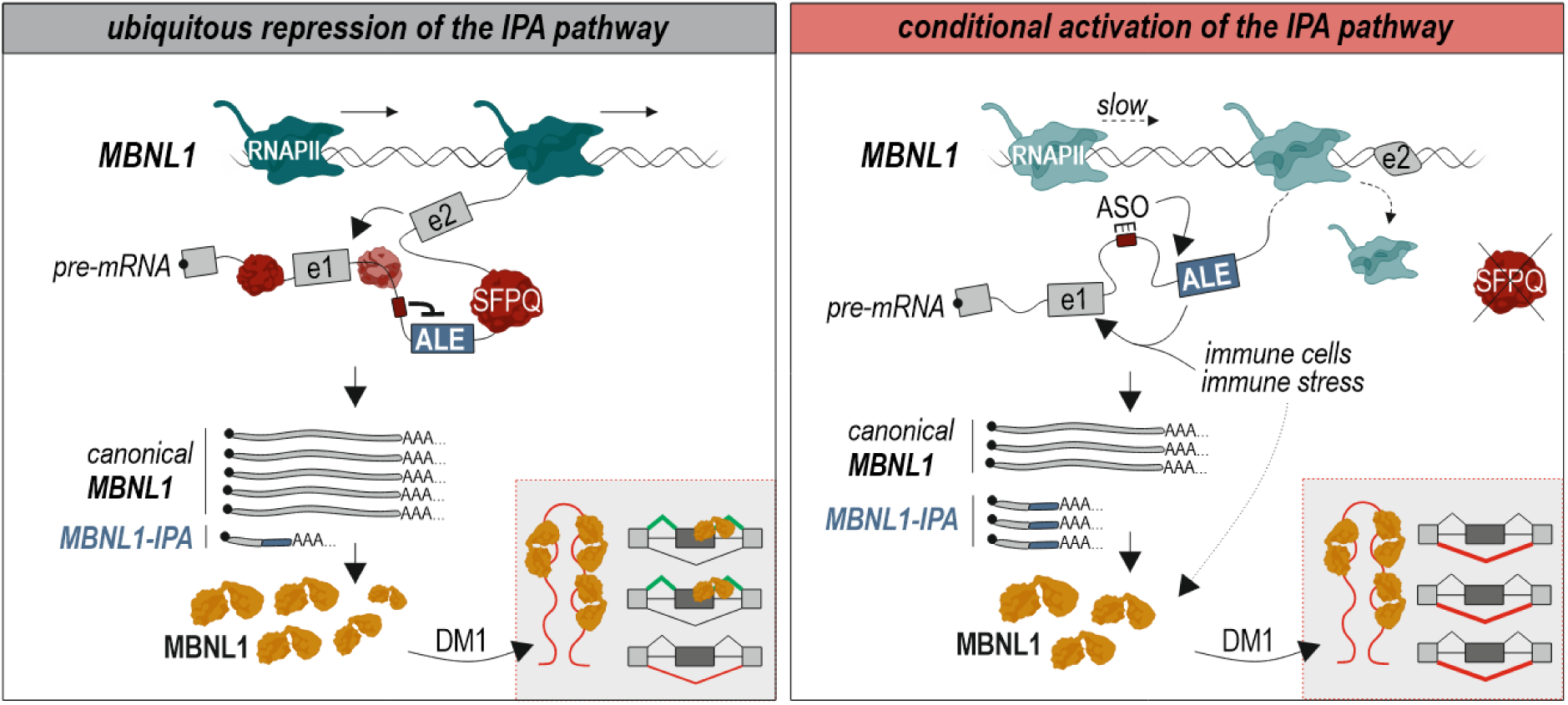
Model of IPA-mediated regulation of *MBNL1* expression. Under basal conditions, productive transcription through the long intron 1 of *MBNL1* is maintained by efficient RNAPII elongation kinetics, SFPQ-dependent preservation of intron integrity, and a cis-regulatory element that suppresses recognition of the cryptic ALE-associated 3′ss. Together, these mechanisms favor canonical splicing and production of full-length *MBNL1* transcript. In immune cells and under conditions that compromise long-intron processing, including innate immune stress, reduced RNAPII elongation rate, SFPQ depletion, or disruption of the cis-regulatory element, transcript processing is redirected toward the IPA pathway. Increased ALE utilization promotes generation of the *MBNL1-IPA* isoform at the expense of canonical *MBNL1* transcript, contributing to reduced MBNL1 abundance and altered the MBNL-dependent alternative splicing pattern, leading to exacerbation of the DM1 molecular phenotype.

Previous studies have shown that IPA often generates truncated transcripts while simultaneously reducing synthesis of full-length gene products, particularly in genes characterized by long introns and complex transcriptional landscapes^7, 9, 11, 14, 16, 18–20, 65^. Our findings place *MBNL1* within this growing class of IPA-regulated genes and provide an example of how IPA can control the abundance of dosage-sensitive RNA-processing factors. Notably, *MBNL1-IPA* is not merely a transient by-product of premature transcription termination but a stable polyadenylated transcript characterized by defined cleavage and splicing patterns. Furthermore, independent lines of evidence, including ribosome profiling, polysome fractionation, and epitope-tagging experiments, support its engagement with the translational machinery. Whether *MBNL1-IPA* is translated into an endogenous microprotein with biological activity remains an important question for future studies.

The repression of the *MBNL1* IPA pathway relies on multiple regulatory layers acting across the exceptionally long intron 1 of *MBNL1*. Among the factors controlling the *MBNL1* IPA pathway, RNAPII elongation kinetics emerged as an important determinant. We show that slower RNAPII elongation favors utilization of the *MBNL1* IPA pathway and attenuates production of canonical *MBNL1* isoforms, consistent with previous studies linking slower RNAPII elongation to increased usage of weak splice sites and intronic polyadenylation signals^14, 18, 53^. Because transcription elongation kinetics vary across genes, tissues, developmental stages, and physiological conditions^66^, modulation of RNAPII dynamics may represent an important mechanism controlling MBNL1 dosage in response to changing cellular states. Furthermore, SFPQ emerged as a particularly prominent regulator, consistent with previous studies identifying SFPQ as key regulator of long-gene expression and a suppressor of cryptic intronic RNA-processing events^59, 60, 69,70^. The strong dependence of *MBNL1* on SFPQ suggests that *MBNL1* expression may be compromised in conditions that perturb SFPQ function, including cellular stress and neurodegenerative disorders such as ALS^15, 67^, Alzheimer and frontotemporal dementia^68^. We further identified a potent cis-regulatory element that acts as a major local determinant of canonical *MBNL1* expression. This regulatory element may provide an additional avenue for therapeutic modulation of *MBNL1* expression. This may be particularly relevant in malignancies in which MBNL1 contributes to disease progression, including MLL- and KMT2A-rearranged leukemias, where reducing MBNL1 abundance has been proposed as a potential therapeutic strategy^42, 44^. Conversely, inhibition of ALE inclusion could represent a means of increasing endogenous MBNL1 expression in disorders associated with insufficient MBNL activity. The integration of these regulatory mechanisms may provide robustness under basal conditions while preserving the capacity for rapid shifts in transcript output when transcriptional or RNA-processing programs are remodeled.

Another important finding of this study is that the *MBNL1* IPA pathway remains functionally relevant even under basal conditions, when ALE usage is relatively low in most non-immune cells. Genetic removal of the ALE region increased canonical *MBNL1* expression and enhanced MBNL-dependent splicing despite producing only modest changes in MBNL1 protein abundance. The physiological significance of the IPA pathway became even more apparent under innate immune stress conditions. Poly(I:C)-induced activation of antiviral signaling promoted *MBNL1-IPA* expression, reduced MBNL1 protein abundance, and altered transcript output from the *MBNL1* locus. Importantly, deletion of the ALE region largely abolished stress-induced reduction of *MBNL1* expression, demonstrating that ALE contributes directly to this response. Because innate immune activation induces widespread remodeling of transcription and RNA processing, additional mechanisms likely cooperate with the IPA pathway to regulate MBNL homeostasis^61, 69–71^. Nevertheless, our findings identify *MBNL1* IPA pathway as a previously unappreciated component of the cellular stress response that contributes to dynamic regulation of *MBNL1* expression. These observations may be particularly relevant in the context of neurodegenerative disorders. In FECD and DM, where functional MBNL levels are already limiting due to sequestration by expanded repeat-containing transcripts, even transient reduction in protein abundance may have disproportionate effects on downstream RNA-processing. Consistent with this possibility, poly(I:C)-induced innate immune signaling reduced MBNL1 protein abundance and worsened the MBNL-dependent AS pattern in DM1-derived fibroblasts. Although the magnitude of these changes was modest, the nonlinear relationship between MBNL dosage and splicing regulation suggests that relatively small fluctuations in MBNL levels may have biologically meaningful consequences. Our findings therefore raise the question of whether other stress conditions associated with increased IPA activation and transcriptional remodeling including viral infection^72, 73^, chronic inflammation and aging^74^, cancer therapies^11^ may act as disease modifiers in disorders characterized by limiting MBNL activity. Future studies will be required to determine whether activation of the *MBNL1* IPA pathway occurs in DM affected tissues in vivo and whether it contributes to clinical variability among patients.

## MATERIALS and METHODS

### Genetic construct preparation

All plasmids were generated using CloneAmp DNA Polymerase (Takara) for PCR amplification and NEBuilder HiFi DNA Assembly Master Mix (NEB) for DNA assembly according to the manufacturers’ protocols. To generate the e1-ALE minigene reporter, a 2826-bp genomic fragment encompassing *MBNL1* exon 1 and the intronic region containing the ALE was amplified using primers MB1InsertF and MB1InsertR. The PCR product was assembled into EcoRI/XhoI-linearized 3×FLAG-pcDNA3.1. The e1-ALE-e2 minigene reporter was generated by amplifying a 1231-bp genomic fragment containing *MBNL1* exon 2 together with approximately 1 kb of upstream intronic sequence using primers I1-1000-e2F3 and I1-1000-e2R3. The resulting fragment was inserted into the XhoI-linearized e1-ALE construct by HiFi assembly. To generate the 101mt mutant reporters, a synthetic annealed oligonucleotide duplex (101GCmt-oligoF/101GCmt-oligoR) carrying the desired nucleotide substitutions was inserted into e1-ALE and e1-ALE-e2 plasmids linearized by PCR using primers 101GCmt2-vec-F and 101GCmt2-vec-R. The 101Δ reporters were generated by inverse PCR using primers 101-GC-deltaF and 101-GC-deltaR followed by plasmid recircularization. For CRISPR-Cas6 genome editing, sgRNAs were designed using CRISPOR and cloned into pSpCas6(BB)-2A-Puro (PX456; Addgene #62688). Individual sgRNA plasmids were generated by inserting annealed oligonucleotides into BbsI-HF-digested PX456 as previously described^75^. Control sgRNAs were adopted from published studies^76, 77^. To generate dual-sgRNA constructs, a second U6-sgRNA expression cassette was introduced into PX456 already carrying the first sgRNA. Briefly, the recipient plasmid was linearized with XbaI, and a complete sgRNA expression cassette containing the second guide sequence was amplified using GoTaq DNA Polymerase (Promega) and primers gRNA-2nd-PX458i6_F and gRNA-2nd-P458i6_2R. The amplified cassette was inserted by HiFi assembly. Simultaneous cleavage by both sgRNAs was designed to generate an approximately 834-bp genomic deletion encompassing the ALE-associated 3′ splice site and adjacent regulatory sequences. pcDNA5/FRT/TO-Rbp1-WT-Am^R^ and pcDNA5/FRT/TO-Rbp1-R746H-Am^R^ were a kind gift of David Bentley^78^. The pEGFP-MBNL1-41 expression vector has also been described previously^40^. All sgRNA sequences and cloning primers are listed in Supplementary Table S1. All constructs were verified by Sanger sequencing prior to use.

### Cell culture and transfection

HeLa and HEK263 cells (ATCC) were maintained in high-glucose DMEM (Lonza) supplemented with 10% foetal bovine serum (FBS; Sigma-Aldrich) and penicillin-streptomycin (Sigma-Aldrich). THP1 cells were cultured in RPMI-1640 medium (CytoGen) supplemented with 10% FBS and penicillin-streptomycin. GM03132 DM1 patient-derived fibroblasts (Coriell Institute) were maintained in Minimum Essential Medium (MEM) supplemented with 10% FBS, non-essential amino acids, and penicillin-streptomycin. All cell lines were cultured at 37°C in a humidified atmosphere containing 5% CO_2_. For transient transfection, cells were seeded in 6-, 12-, or 24-well plates and transfected at approximately 50–60% confluence. Plasmid DNA was delivered using Lipofectamine 3000 (Thermo Fisher Scientific), whereas siRNAs and antisense oligonucleotides (ASOs) were transfected using Lipofectamine RNAiMAX (Thermo Fisher Scientific) according to the manufacturer’s instructions. *MBNL1* was depleted using an siRNA targeting the 3′UTR of *MBNL1*^76^ (Sigma Aldrich). *SFPǪ* knockdown was performed using either a single siRNA (s12712; Ambion) or a pool of four siRNAs (sc-38304; Santa Cruz Biotechnology). A previously described non-targeting siRNA was used as a control^80^ (Sigma Aldrich). Unless stated otherwise, siRNAs were used at a final concentration of 50 nM, and cells were harvested 48 h after transfection. ASOs carrying 2′-O-methoxyethyl-phosphorothioate (2′MOE-PS) modifications (Polgen) were used at a final concentration of 100 nM. Cells were harvested 48 h after transfection. For co-treatment experiments, cells were first transfected with siRNAs (siCtrl or siSFPQ #2) or ASOs (ASO-Ctrl or ASO-101). After 24 h, minigene reporter plasmids (150 ng per well) were introduced using Lipofectamine 3000. Cells were harvested 24 h later for RNA and protein analyses. Sequences of all siRNAs, ASOs, and oligonucleotides used in this study are provided in Supplementary Table S1.

### Generation of ALE knockout cell lines by CRISPR-CasG genome editing

To identify the most efficient sgRNAs for deletion of the ALE region, individual sgRNAs were first evaluated in HEK263 cells. Cells were transfected with PX456 plasmids expressing a single sgRNA, and genomic DNA was isolated 48 h post-transfection using QuickExtract™ DNA Extraction Solution 1.0 (Lucigen). The targeted loci were amplified using GoTaq DNA Polymerase (Promega) and primers ALE_gRNAtest1_F and ALE_gRNAtest1_R. PCR products were analyzed by agarose gel electrophoresis, purified, and subjected to Sanger sequencing. Genome editing efficiencies were quantified using TIDE and Synthego ICE. Based on these analyses, the two most efficient sgRNAs were selected to generate a dual-guide construct. One sgRNA expression cassette was amplified using the gRNA-2nd-PX458i6_F and gRNA-2nd-PX458i6_R primers and inserted into an XbaI-linearized PX458i6 vector harboring the second sgRNA expression cassette. To generate ALE knockout cell lines, a monoclonal parental HEK263 cell line was transfected with the dual-sgRNA PX456 construct using Lipofectamine 3000 (Thermo Fisher Scientific). 48 h after transfection, cells were seeded at low density and subjected to puromycin selection (1 μg/mL) for an additional 48 h. Following selection, cells were maintained in puromycin-free growth medium to allow recovery and clonal expansion. Surviving cells were subsequently replated onto 15-cm dishes, and individual colonies were isolated using Pyrex® cloning cylinder (Merck) and expanded for genotyping. For clone screening, genomic DNA was isolated and the targeted region was analysed by PCR using primers F1 and R1, which flank the expected deletion. To distinguish homozygous deletions, an additional PCR assay was performed using primers F1 and R2, with one primer positioned within the deleted region. Candidate homozygous clone was further validated by Sanger sequencing.

### Stress induction

To identify stress conditions the *MBNL1* IPA pathway is sensitive to, HeLa cells were exposed to polyinosinic:polycytidylic acid (poly(I:C); 0.5 µg/mL; InvivoGen), hydrogen peroxide (H₂O₂; 50µM; Merck) or thapsigargin (TG; 3µM; Merck) for 6 h, or sodium arsenite (NaAsO_2_; 0.1µM) (Merck) for 45min followed by 2 h of post-stress induction (PtSI) period in fresh medium. Poly(I:C) was delivered using Lipofectamine RNAiMAX (Thermo Fisher Scientific) according to the manufacturer’s instructions. Control samples were treated with the corresponding vehicles: RNAiMAX for poly(I:C), DMSO for TG, H_2_O for H₂O₂ and NaAsO_2_ treatment. Transcript abundance was quantified by RT-qPCR using *RPLP0* as a reference gene for poly(I:C), *ACTB* for H₂O₂, *GAPDH* for TG and NaAsO_2_ treatment. For extended poly(I:C) experiments, WT (c4) and ALE KO (c10^-/-^) HEK263 were treated with poly(I:C) at a final concentration of 0.5 μg/mL for 6 h. GM03132 fibroblasts were treated with 0.1 μg/mL poly(I:C) under identical conditions. Cells were harvested either immediately after treatment for RNA analysis or following an additional 18 h PtSI period for RNA and protein analyses. All H₂O₂ treatments were performed in serum-free medium.

### Poly(A)-tail length assay

Poly(A)-tail length was analysed using the Poly(A) Tail-Length Assay Kit (Thermo Fisher Scientific) according to the manufacturer’s instructions. Briefly, 2 μg of total RNA isolated from THP1 cells was subjected to G/I tailing, followed by reverse transcription, including a no-reverse-transcriptase (RT-) control reaction. The resulting cDNA was amplified using a tail-specific reverse primer and the MBNL1-specific forward primer MBALEF (5′-TGGAGCTTCATACCCAGG-3′). The expected amplicon size excluding the poly(A) tail was 427 bp. PCR products were resolved by agarose gel electrophoresis, and fragments ranging from 400 to 1000 bp were excised and purified. Purified DNA was subjected to Sanger sequencing, and sequencing chromatograms were analysed using VectorBee software to estimate poly(A)-tail length. To confirm the identity of the amplified transcript, PCR products were additionally analysed using the MBNL1-IPA-specific primer pair 3RACEgSPECoutF (5′-GGCCACAGAACAGTCCCTTT-3′) and MBNL1_ALER3 (5′-AGTTTGCATTGATTAGCTGCTGT-3′), which generates a 161-bp amplicon.

### RNA stability assay

RNA stability was assessed following transcriptional arrest with actinomycin D (ActD; Merck). HEK263 cells were seeded in 12-well plates and treated with ActD at a final concentration of 5 µg/mL. Cells were harvested at the time points indicated in the corresponding figure, and total RNA was isolated using TRI Reagent™ Solution (Thermo Fisher Scientific) according to the manufacturer’s instructions. Gene expression levels were quantified by RT-qPCR, normalized to *GAPDH*, and expressed relative to the corresponding transcript level at time point -T0 prior to ActD treatment.

### Splicing and gene expression analyses

Total RNA was isolated from cultured cells and from a pellet of CD14^+^ monocytes (hMoCD14^+^; Merck) using TRI Reagent (Sigma-Aldrich), followed by purification with the Total RNA Zol-Out Kit (AsA Biotechnology), including on-column DNase treatment according to the manufacturer’s instructions. First-strand cDNA was synthesized from 1-2 μg of total RNA using the High-Capacity cDNA Reverse Transcription Kit (Thermo Fisher Scientific). For alternative splicing analyses, PCR amplification was performed using GoTaq® Flexi DNA Polymerase (Promega). PCR products were separated on agarose gels containing ethidium bromide (Merck) and visualized using a G:BOX imaging system (Syngene). Band intensities were quantified using Multi Gauge v3.0 software. Alternative splicing was quantified as percent spliced in (PSI). Primers were designed to amplify across a single alternative splicing event, and the size difference between alternative amplicons was maintained below 200 nt to minimize amplification and quantification bias. Gene expression analyses were performed by quantitative PCR (RT-qPCR) using Maxima SYBR Green/ROX Master Mix (Thermo Fisher Scientific) on QuantStudio™ 6 and QuantStudio™ 7 Flex Real-Time PCR Systems (Thermo Fisher Scientific) according to the manufacturer’s instructions. Primer sequences used in this study are provided in Supplementary Table S1.

### DRB-based analysis of transcription

Transcriptional elongation was transiently inhibited using 5,6-dichloro-1-β-D-ribofuranosylbenzimidazole (DRB; Merck). HeLa cells were seeded in 6-well plates and treated with either DMSO (control) or DRB at a final concentration of 100 µM for 4 h. Following treatment, cells were washed twice with PBS and released in fresh culture medium. Cells were harvested at the time points indicated in the corresponding figure over a 65 min time course. Total RNA was isolated using TRI Reagent (Sigma-Aldrich) and treated with DNase prior to cDNA synthesis using the High-Capacity cDNA Reverse Transcription Kit (Thermo Fisher Scientific). Transcript level was quantified by RT-qPCR using primers specific for regions spanning the *MBNL1* pre-mRNA. Expression level at each time point post DRB-treatment was referenced and normalized to the corresponding transcript level at time point T0 post DMSO-treatment. Primer sequences used in this assay are provided in Supplementary Table S1.

### RNA Polymerase II elongation assay

HEK263 cells were seeded in 12-well plates and transfected with 2 µg of expressing plasmid with either an α-amanitin-resistant wild-type *POLR2A* catalytic subunit of RNA polymerase II (RNAPII am^R^ WT) or an α-amanitin-resistant slow-elongation mutant carrying R746H substitution (RNAPII am^R^ slow)^78^. Both constructs contain a mutation within a sequence encoding the funnel domain of RNAPII that confers resistance to α-amanitin. 16 h after transfection, cells were treated with α-amanitin (Merck) at a final concentration of 2.5 µg/mL to inhibit endogenous RNA polymerase II activity while preserving transcription driven by the exogenous α-amanitin-resistant RNAPII proteins. Non-transfected cells cultured in the presence or absence of α-amanitin served as controls. Cells were harvested 48 h after transfection for RNA and protein analyses.

### Immunoblotting

Cells were lysed in RIPA buffer (150 mM NaCl, 50 mM Tris-HCl pH 8.0, 1 mM EDTA, 0.5% NP-40, 0.5% Triton X-100, 0.5% sodium deoxycholate, and 0.1% SDS) supplemented with Halt™ EDTA-free Protease Inhibitor Cocktail (Thermo Fisher Scientific). Lysates were incubated on ice for 30 min, snap-frozen at −80°C overnight, and clarified by centrifugation at 18,000 x g for 15 min at 4°C. Protein concentrations were determined using the Pierce BCA Protein Assay Kit (Thermo Fisher Scientific). Equal amounts of protein (10-15 µg) were mixed with Bolt™ reducing agent and Bolt™ LDS sample buffer (Thermo Fisher Scientific), incubated at 70°C for 10 min, and separated on Bolt™ 4–12% Bis-Tris Plus gradient gels (Thermo Fisher Scientific). Proteins were transferred 0.45-µm PVDF membranes (Sigma-Aldrich). Wet transfer was performed at 100 V for 1 h at 4°C, except for His-tagged RNAPII, which was transferred at 20 V overnight at 4°C. Membranes were blocked for 1 h in 5% (w/v) skim milk prepared in TBST (Tris-buffered saline supplemented with 0.1% Tween-20) and incubated with primary antibodies listed in Supplementary Table S1. For SFPQ detection, membranes were processed using SNAP i.d. system (Merck) in TBST containing 1% BSA. HRP-conjugated secondary antibodies were used where appropriate. Immunoreactive signals were detected using Immobilon™ Forte Western HRP substrate (Sigma-Aldrich) and visualized using a ChemiDoc™ imaging system (Bio-Rad).

### RNA immunoprecipitation

RNA immunoprecipitation (RNA-IP) was performed using HeLa cells. Cells were lysed in RIPA buffer supplemented with Halt™ EDTA-free Protease Inhibitor Cocktail (Thermo Fisher Scientific), 1 mM DTT, and 40 U RNasin® (Promega). Lysates were incubated on ice for 30 min, snap-frozen at −80°C overnight, and homogenized by five passages through a 25-gauge needle. Following clarification by centrifugation at 2,000 rpm for 10 min at 4°C, lysates were incubated with 1 µg of anti-SFPQ antibodies (ab-SFPQ, ab177146, Abcam; or ab-SFPQ-2, ab66357, Abcam) or rabbit IgG control antibody (ab37415, Merck) for 1 h at 4°C on a rotating wheel. Antibody-lysate complexes were incubated with Dynabeads™ Protein A (Invitrogen) for 2 h at 4°C with rotation. Beads were washed three times with PBST buffer (137 mM NaCl, 2.7 mM KCl, 8 mM Na₂HPO₄, 2 mM KH₂PO₄, 0.02% Tween-20), and bound complexes were eluted using acidic glycine solution. RNA was isolated using the PureLink™ RNA Mini Kit (Thermo Fisher Scientific) according to the manufacturer’s instructions. RIP efficiency was quantified by RT–qPCR and expressed as the relative IP/Input ratio, calculated according to the following formula: 2^((CtIP−CtInput)−(CtIgG−CtInput,IgG))^. Primer sequences are provided in Supplementary Table S1.

### RNA-seq data analysis and Sashimi plot visualization

Poly(A)-selected RNA-seq datasets were obtained from the Gene Expression Omnibus (GEO) for HepG2 cells (GSE88362)^81^, CD14^+^ monocytes (GSE60272), PBMC (GSE58335)^82^ and B-cells (GSE60268)^81^. Sequencing reads were aligned to the human reference genome (GRCh38) using the STAR v2.7.6a^83^. Sashimi plots were generated using ggsashimi v1.1.5^84^. Two regions of the *MBNL1* locus were visualized: the e1–ALE/e2 region (chr3: 152,266,405-152,415,265) and the e1-ALE region (chr3: 152,266,405-152,302,666). Gene models were based on the Ensembl GRCh38 annotation. PAS-seq data from HepG2 (GSE218677)^51^ were downloaded in BigWig format from GEO and visualized using the University of California, Santa Cruz (UCSC) Genome Browser. To examine RNA processing within the intron 1 of *MBNL1*, poly(A)+ RNA-seq datasets generated by the ENCODE Consortium were analyzed through the ENCODE RNA-seq supertrack in the UCSC Genome Browser^85^. Representative datasets used for visualization in Figure 1A included PBMC (ENCSR406JSP/ENCFF725SYE), B-cells (ENCSR000CTV/ENCFF806CLT), CD14^+^ monocytes (ENCSR000CUC/ENCFF665WHG), spleen (ENCSR610Q0X/ENCFF508VHM), thymus (ENCSR775KCE/ENCFF378LKK), brain (ENCSR274JRR/ENCFF263GSB), heart (ENCSR635GTY/ENCFF484GAK), skeletal muscle myoblasts (ENCSR000COY/ENCFF680GTA), HepG2 cells (ENCSR561FEE/ENCFF266YJG) and HeLa cells (ENCSR000CPR/ENCFF615XWV). To assess the effects of RNA-binding protein depletion on *MBNL1* transcript processing, ENCODE RNA-seq datasets generated following shRNA-mediated knockdown of selected RBPs in HepG2 cells were analyzed^85^. Datasets visualized in Figure 4A included sh-*SFPǪ* (ENCSR782MXN/ ENCFF464QJC), sh-*hnRNPM* (ENCSR665JMS/ENCFF226GYG), sh-*hnRNPC* (ENCSR052IYH/ ENCFF588YAG), and sh-*PTBP1* (ENCSR064DXG/ENCFF776JVF). To evaluate SFPQ occupancy across the *MBNL1* locus, ENCODE eCLIP-seq datasets^85^ were examined. A representative HepG2 dataset (ENCSR665DLL/ENCFF681EDG) was selected for visualization in Figure 4B.

### Splice-site strength prediction

Splice-site strength was evaluated using the MaxEntScan algorithm^54^, which predicts splice-site recognition potential based on maximum entropy modeling. Sequences encompassing the ALE-associated 3′ ss were analyzed using the MaxEntScan implementation available through GeneBe. Predicted splice-site strength was expressed as the MaxEntScan score, with higher values indicating greater similarity to consensus splice-site features.

### Statistical analysis

Statistical analyses were performed using the GraphPad Prism 10 (license GPA-2667082-RJVN-2ACA4). Unless otherwise indicated, experiments were conducted using at least three independent biological replicates (*N*), and data are presented as mean± standard deviation (SD). Statistical significance was assessed using two-tailed Student’s *t*-tests. For experiments involving multiple comparisons, *P* values were adjusted using the Holm-Sidak method. Statistical significance was defined as *P* < 0.05 and is indicated as follows: *P* < 0.05 (∗), *P* < 0.01 (∗∗), and *P* < 0.001 (∗∗∗), *P* > 0.05 (ns),

## DATA AVAILABILITY STATEMENT

All data supporting the findings of this study are included in the article and its Supplementary Information. Additional information is available from the corresponding authors upon reasonable request.

## AUTHOR STATEMENT

KT, KS: conceptualization; KT, KS: funding acquisition; KT, KK, MJ, EK, AB, DH, ŁJS: data collection; KT, methodology; KT: figure preparation; KT: writing; ŁJS, KS, MJ, KK, DCHE, AB, EK, MMM: editing.

## ACKNOWLEDGEMENTS

This work was supported by the National Science Centre, Poland (2020/39/D/NZ3/01658 to KT and 2020/38/A/NZ3/00498 to KS), Myotonic Dystrophy Foundation High Priority Short-Term Project Award (to KT), and Early Career Scholar Award (to KT), Foundation for Polish Science co-financed by the European Union under the European Funds for Smart Economy 2021-2027 (FENG.02.01-IP.05-M038/25 to KS). The authors thank Agata Stępień for valuable discussions and insights into 3′-end RNA sequencing data, and Agnieszka Piasecka for helpful discussions and critical feedback on the manuscript. The authors acknowledge the use of Microsoft Copilot (Microsoft Corporation) for assistance with language editing, grammar correction, and stylistic refinement of the manuscript. All AI-assisted outputs were reviewed, edited, and verified by the authors, who take full responsibility for the final manuscript and its scientific content.

## Notes

### Competing Interest Statement

The authors have declared no competing interest.

