## Supplemental Figures and Methods for "Dynamic regulation of MBNL1 dosage through a stress-responsive intronic polyadenylation pathway"

#### SUPPLEMENTARY FIGURES

**Supplementary Figure S1.** Evidence for an active immune cell-enriched IPA pathway in the *MBNL1* locus.

**Supplementary Figure S2.** Translational potential of *MBNL1-IPA* RNA.

**Supplementary Figure S3.** MBNL-dependent exon 1 splicing and transcription kinetics contribute to *MBNL1-IPA* expression.

**Supplementary Figure S4.** SFPQ suppresses activation of the *MBNL1* IPA pathway.

**Supplementary Figure S5.** A potent intronic cis-regulatory element represses the *MBNL1* IPA pathway.

**Supplementary Figure S6.** Deletion of the ALE region increases *MBNL1* expression and enhances the MBNL-dependent splicing pattern.

**Supplementary Figure S7.** Poly(I:C)-induced innate immune stress remodels *MBNL2* transcript output and reduces MBNL2 protein abundance.

**Supplementary Figure S8.** Poly(I:C)-induced innate immune stress reduces MBNL1 abundance and impairs MBNL-dependent splicing in DM1-affected fibroblasts.

### Figure S1

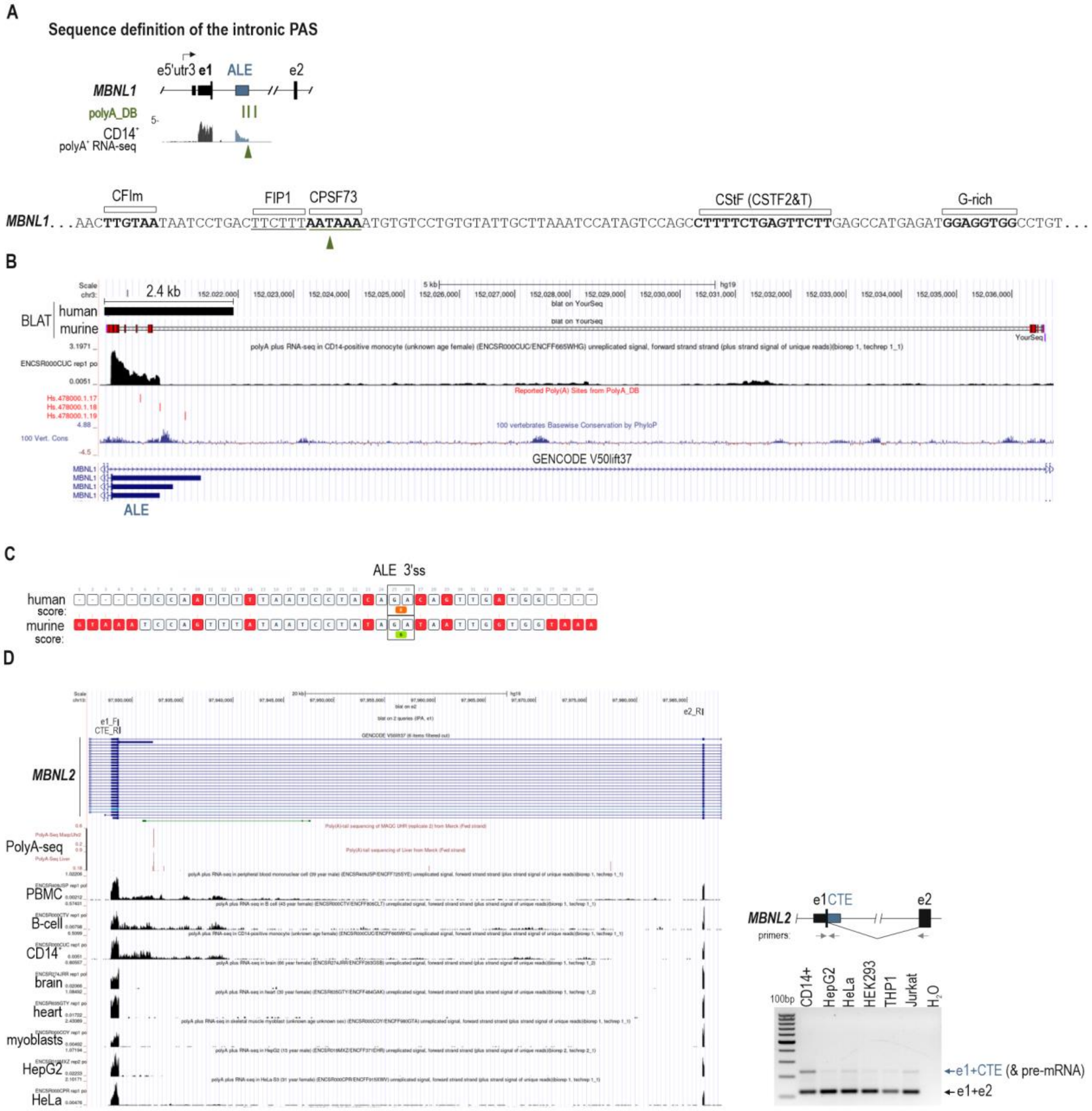
A
Sequence definition of the intronic PAS
e5'utr3 e1
ALE
e2
MBNL1
polyA_DB
CD14+
polyA+ RNA-seq
CFIm
FIP1
CPSF73
CStF (CSTF2&T)
G-rich
MBNL1. . . AACTTGTAATAATCCTGACTTCTTTAATAAAATGTGTCCTGTGTATTGCTTAAATCCATAGTCCAGCCTTTTCTGAGTTCTTGAGCCATGAGATGGAGGTGGCCTGT . . .
B
2.4 kb
BLAT
human
murine
GENCODE V50lift37
ALE
C
ALE 3'ss
human
score:
murine
score:
D
MBNL2
PolyA-seq
PBMC
B-cell
CD14+
brain
heart
myoblasts
HepG2
HeLa
e1CTE
e2
primers:
100bp
CD14+
HepG2
HeLa
HEK293
THP1
Jurkat
H2O
e1+CTE (& pre-mRNA)
e1+e2

#### Figure S1 continuation

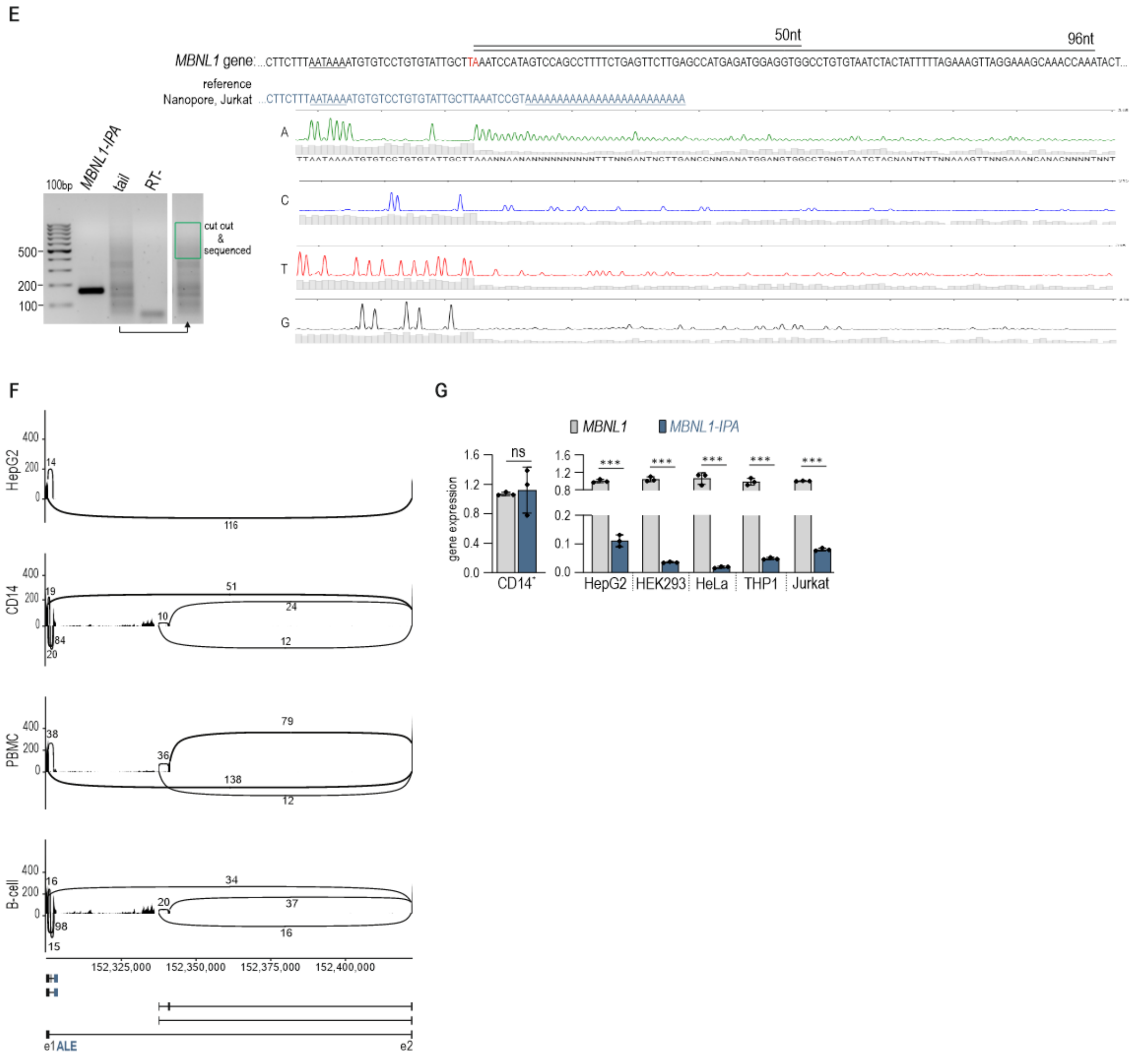


#### Supplementary Figure S1. Evidence for an active immune cell-enriched IPA pathway in the *MBNL1* locus.

A) Schematic representation of the *MBNL1* locus encompassing e1 – ALE region. The position of the exonized intronic sequence (ALE; blue), annotated polyadenylation sites from PolyA_DB (green), and representative poly(A)+ RNA-seq coverage from CD14⁺ monocytes from ENCODE database are shown. The lower panel displays the nucleotide sequence surrounding the intronic PAS. Predicted binding sites for cleavage and polyadenylation-associated factors including CFIm (CPSF4), FIP1, CPSF73-containing CPSF complex, and CstF (CSTF2/T), as well as a G-rich downstream sequence element, are indicated. The canonical polyadenylation signal (AATAAA) is highlighted in green and marked with an arrow.

B) UCSC Genome Browser view of the human *MBNL1* ALE sequence and surrounding intron 1. A 2.4 kb-long murine sequence encompassing a region potentially corresponding to the human ALE sequence was aligned to the human genome using BLAT tool, together with the corresponding 2.4-kb human sequence used as a reference. The top track indicates a fully aligned 2.4kb-long human sequence (black box), and partially aligned 2.4-kb murine sequence (red boxes). The remaining tracks show poly(A)+ RNA-seq coverage from CD14⁺ monocytes, PolyA_DB-annotated polyadenylation sites, vertebrate PhyloP conservation coverage, and

GENCODE transcript annotations. The region corresponding to the human ALE shows limited conservation at 5’end of ALE and lacks evidence of a corresponding murine intronic PAS.

C) Comparative analysis of the human and murine ALE-associated cryptic 3'ss using online splice-site prediction algorithm MaxEntScan[1]. The strength of human and murine splice acceptor site is indicated in orange and green boxes below nucleotides. Common nucleotides are marked in white boxes, nucleotides that differ between human and mouse sequence are marked in red boxes.

D) *Left*, UCSC Genome Browser view of a fragment of human *MBNL2* locus showing poly(A)+ RNA-seq ENCODE read coverage across multiple cell types and tissues, together with PolyA_DB annotations and the GENCODE transcripts. A prominent intronic polyadenylation-associated region was detected immediately downstream of *MBNL2* e1 (the first protein coding exon). In contrast to *MBNL1*, the *MBNL2* IPA-associated event incorporates the intervening intronic sequence into e1 prior to intronic PAS generating a composite terminal exon (CTE). Position of primers for three-primer RT-PCR assay is marked as vertical lines. *Right*, schematic representation of the predicted *MBNL2* transcript architecture and RT-PCR analysis using primers located in exon 1 (e1_F), the IPA-associated region (CTE_R), and exon 2 (e2_R). The upper amplicon corresponds to an e1-CTE transcript (together with pre-mRNA/intron retention) and was predominantly detected in CD14$^+$ monocytes, whereas the canonical e1-e2 product was observed in all analyzed cell types. Products were amplified using primers’ ratio 1:1:0.5 (e1_F:CTE_F:e2_R).

E) *Left*, RT-PCR analyses of poly(A)-tail length showing control and A-tail specific amplification of *MBNL1-IPA*. A region corresponding to 400-1000 bp was cut out of the gel (green box) and sequenced. RT-, negative control of reverse transcription indicates no contamination with genomic DNA. *Right*, the reference genomic sequence of *MBNL1* intron 1 and representative nanopore sequencing output derived from Jurkat cells corresponding to the 3’end of *MBNL1-IPA* with a poly(A)-tail. The predicted polyadenylation signal (AATAAA) is highlighted and the putative cleavage site is indicated in red. The corresponding base-calling signals from Sanger sequencing of 400-1000 bp products obtained in poly(A)-tail length assay are shown below.

F) Sashimi plot visualization of alternative cryptic splicing across intron 1 of *MBNL1* in CD14$^+$ monocytes, B cells, PBMCs, and HepG2 cells using ENCODE RNA-seq datasets. Numbers indicate split-read counts supporting individual splice junctions with a cut off > 10 split reads. The schematic representation of the analyzed genomic region is shown below.

G) Comparison of *MBNL1-IPA* expression relative to canonical *MBNL1* transcript level across primary immune cells (CD14$^+$), cancer derived immune cell lines (THP1, Jurkat) and non-immune cell lines (HepG2, HEK293 and HeLa) measured by RT–qPCR. $N = 3$.

**Figure S2**

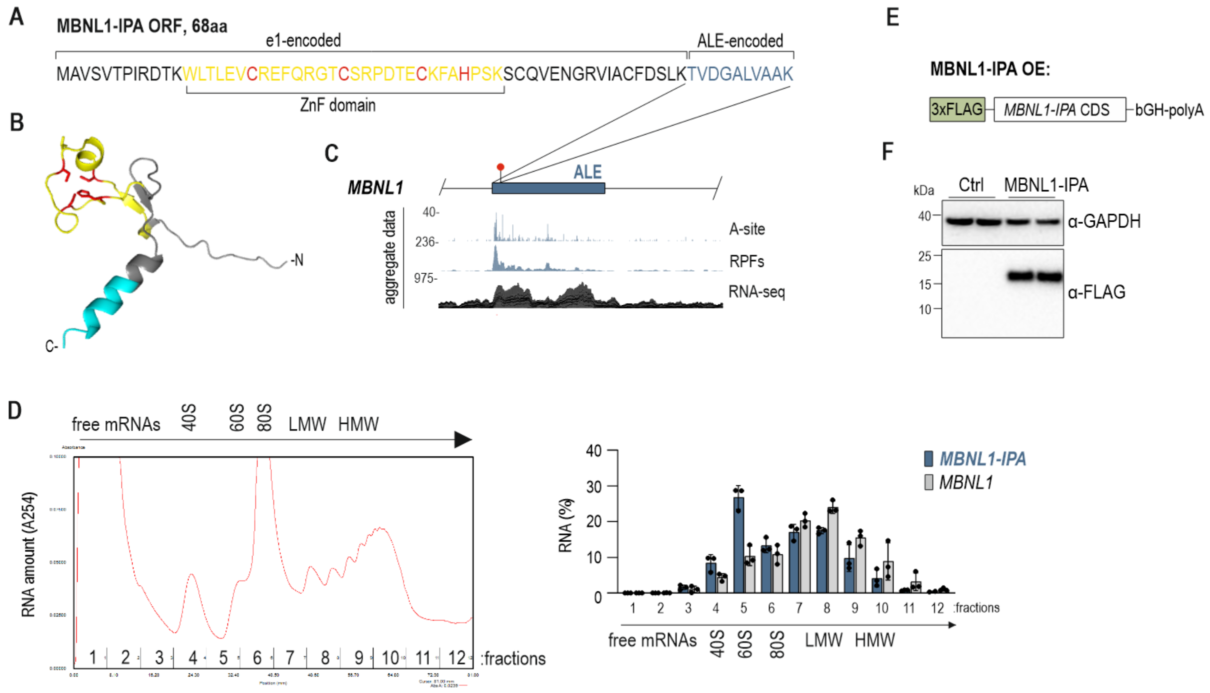


#### Supplementary Figure S2. Translational potential of *MBNL1-IPA* RNA.

A) Predicted open reading frame (ORF) of *MBNL1-IPA*. Regions encoded by exon 1 and ALE are indicated. The predicted CCCH-type zinc-finger (ZnF) domain is highlighted in yellow, and zinc-coordinating residues are shown in red.

B) AlphaFold-predicted structure of the putative *MBNL1-IPA* micropeptide. Structural features are colored as indicated in panel A.

C) Ribosome profiling evidence supporting ribosomal engagement of the *MBNL1-IPA* RNA. Aggregate GWIPS-viz[2] tracks show elongating ribosomes (A-site profiles), ribosome-protected fragments (RPFs), and RNA-seq coverage across the ALE. The predicted translation termination codon is marked by a red dot.

D) Association of *MBNL1-IPA* with ribosomes. *Left*, Representative global RNA polysome profile generated by the linear sucrose gradient. LMW and HMW, light and heavy molecular weight polysomal fractions, respectively. *Right*, the distribution of *MBNL1-IPA* and canonical *MBNL1* transcripts across the sucrose gradient was analyzed by RT-qPCR of RNA isolated from each of the 12 gradient fractions. LMW and HMW indicate light and heavy polysomal fractions, respectively. *N* = 3.

E) Schematic representation of the *MBNL1-IPA* expression construct comprising exon 1-derived and ALE-derived coding sequences fused to a C-terminal 3×FLAG tag.

F) Detection of the MBNL1-IPA-encoded micropeptide by western blotting. HEK293 cells were transfected with the 3xFLAG-MBNL1-IPA expression construct or control vector. The micropeptide was detected using an anti-FLAG antibody. GAPDH served as a loading control.

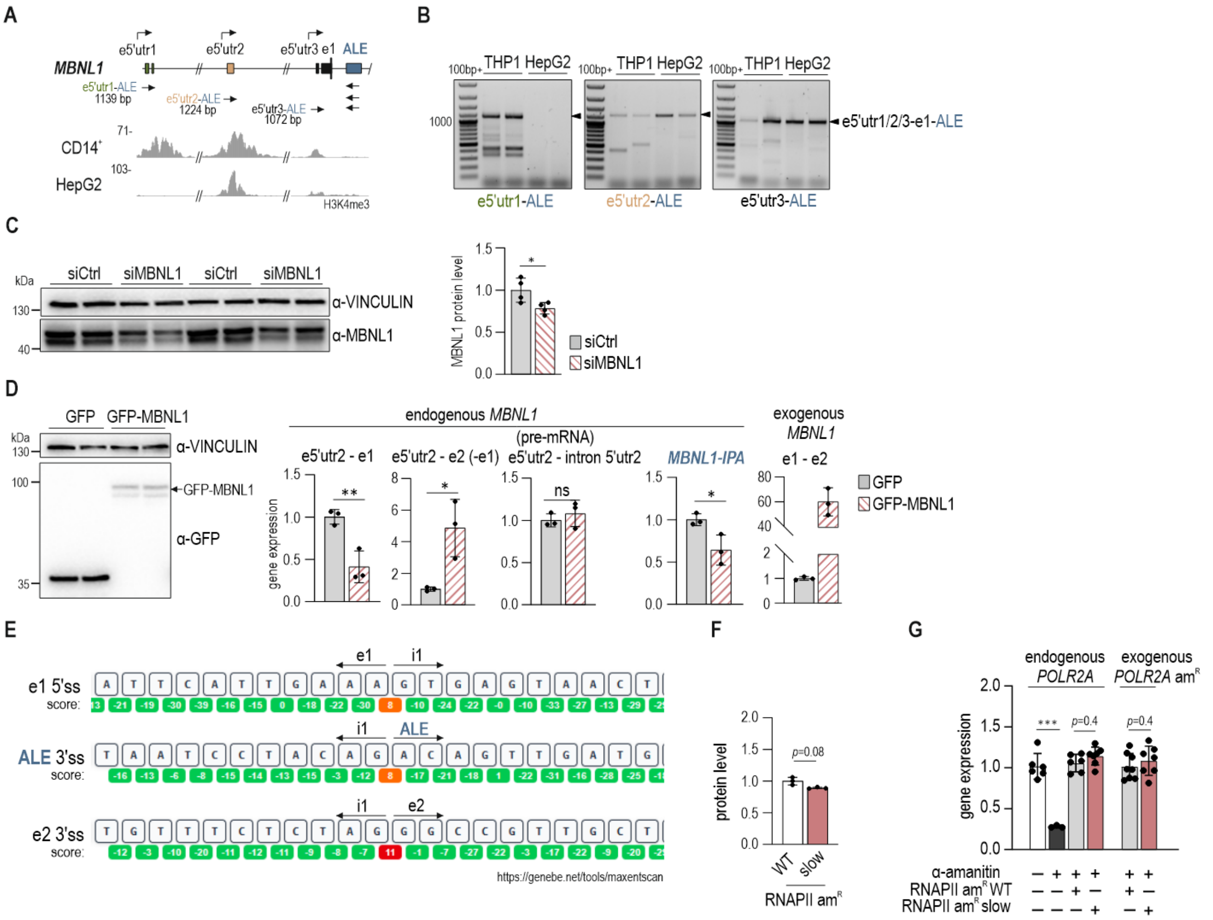


**Supplementary Figure S3. MBNL-dependent exon 1 splicing and transcription kinetics contribute to *MBNL1-IPA* expression.**

A) Genomic organization of the 5' end of the *MBNL1* locus. *Top*, schematic representation of the 5' end of the *MBNL1* locus showing alternative first exons (e5'utr1, e5'utr2, e5'utr3), e1, ALE, locations of RT–PCR primers and the size of amplified products. *Bottom*, representative ENCODE datasets for H3K4me3 ChIP-seq profiles from $CD14^{+}$ monocytes (ENCSR000ASN/ENCFF171ORL) and HepG2 cells (ENCSR000DUF/ENCFF171BSH) displayed across the corresponding genomic region. H3K4me3 enrichment is shown across the promoter regions corresponding to alternative *MBNL1* transcript isoforms.

B) *MBNL1-IPA* is expressed from all three alternative promoters of *MBNL1*. Representative gels showing RT–PCR analysis of *MBNL1-IPA* originating from alternative 5'UTR regions shown in panel A. Primer pairs specific for e5'utr1–ALE, e5'utr2–ALE, and e5'utr3–ALE transcripts were used.

C) HeLa cells were transfected with either siCtrl or siMBNL1 targeting 3'UTR of *MBNL1*. *Left*, western blot analyses showing MBNL1 abundance upon siRNA treatment in Hela cells. VINCULIN served as a loading control. *Right*, quantification of MBNL1 abundance in reference to VINCULIN and normalized to siCtrl-treated cells. *N* = 4.

D) HeLa cells were transfected with either GFP or MBNL1-GFP expression constructs and analyzed by RT-qPCR. *Left*, representative western blot showing GFP-MBNL1 or control GFP overexpression in HeLa cells. VINCULIN served as a loading control. *N* = 4. *Right*, the level of endogenous *MBNL1* isoforms was assessed using primers specific for e1-containing isoforms (e5'utr2-e1), e1-lacking isoforms (e5'utr2-e2, one primer located across a splice junction) and *MBNL1* pre-mRNA showing total level of endogenous *MBNL1* originating from 5'UTR2 (e5'utr2-intron 1). Consistent with the previously described exon 1-mediated autoregulatory feedback mechanism, MBNL1 overexpression reduced the level of endogenous e1-containing isoforms and increased e1-lacking isoforms, while pre-mRNA level remained unchanged. Consistently, *MBNL1-IPA* expression was modestly reduced and followed the characteristic response of the exon 1-dependent autoregulatory pathway. Data are presented relative to *TBP* and normalized to GFP-transfected cells. *N* = 3

E) Computational analysis of splice-site strength within the *MBNL1* region encompassing e1, ALE and e2, performed using MaxEntScan[1]. The canonical exon 1 donor site (e1 5'ss), the cryptic ALE acceptor site (ALE 3'ss), and the canonical exon 2 acceptor site (e2 3'ss) are shown together with adjacent nucleotides. Values beneath individual nucleotides indicate position-specific contributions to splice-site strength. Predicted splice-site scores are highlighted in orange (e1 5'ss and ALE 3'ss) and red (e2 3'ss). Arrows indicate the boundaries of the analyzed splice sites.

F) Quantification of transiently expressed His-tagged α-amanitin-resistant ($am^R$) wild-type (WT) or slow RPB1 subunits in reference to VINCULIN and normalized to WT RPB1. *N* = 3.

G) RT-qPCR analysis of endogenous and exogenous *POLR2A* transcripts in control cells and following α-amanitin treatment. Expression values are shown relative to untreated control cells (for endogenous *POLR2A*) or WT *POLR2A* (for exogenous *POLR2A*). *N* = 6-7.

#### Figure S4

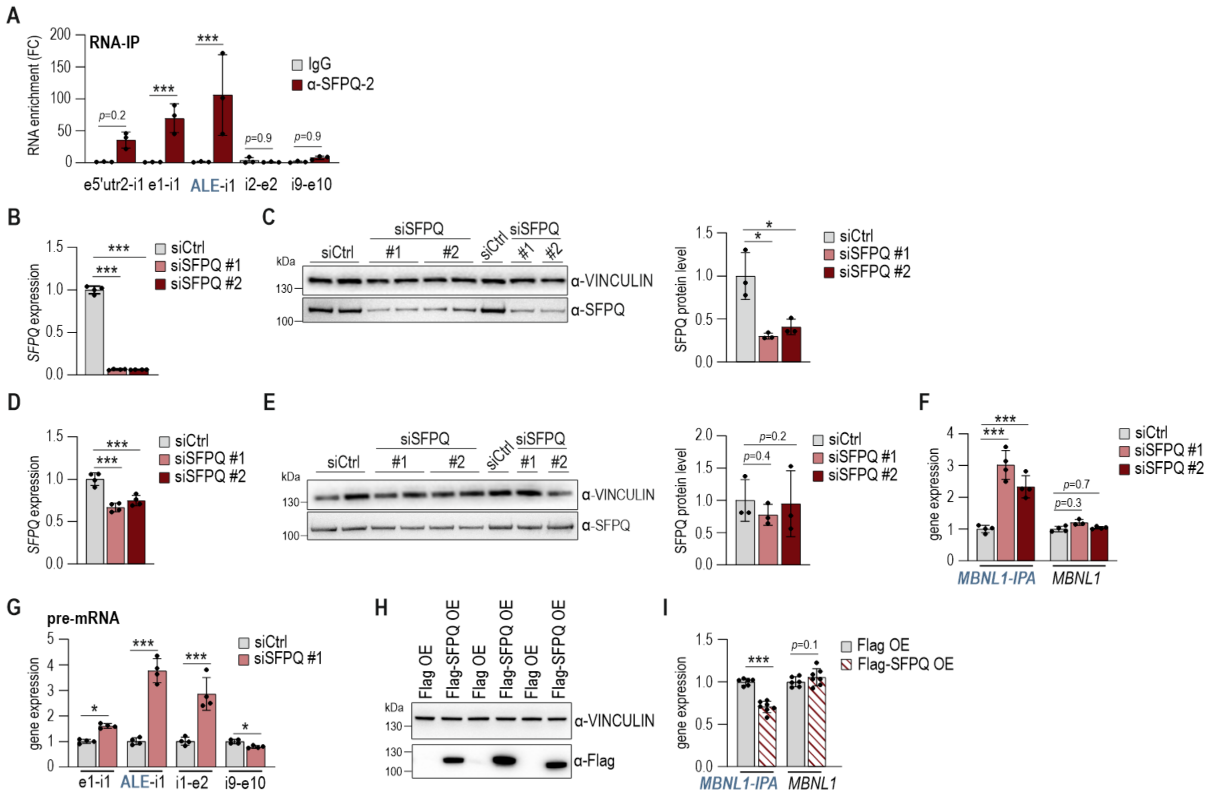

**Supplementary Figure S4. SFPQ suppresses activation of the *MBNL1* IPA pathway.**

A) RNA immunoprecipitation (RNA-IP) performed in HeLa cells using SFPQ antibody (α-SFPQ-2, ab992257) followed by RT-qPCR analysis of the indicated *MBNL1* pre-mRNA regions shown by arrows in the main Figure 4B. RNA enrichment is normalized to the IgG control. *N* = 3.

B) RT-qPCR validation of *SFPQ* depletion in HeLa cells following transfection with siSFPQ #1 (single siRNA) and siSFPQ #2 (siRNAs pool). *SFPQ* expression level is calculated relative to *GAPDH* and normalized to siCtrl-treated cells. *N* = 4.

C) *Left*, western blot analysis of SFPQ protein abundance in HeLa cells following transfection with siSFPQ #1 or siSFPQ #2. VINCULIN served as a loading control. *Right*, quantification of SFPQ abundance in reference to VINCULIN and normalized to siCtrl-treated cells. *N* = 3.

D) RT-qPCR validation of modest *SFPQ* depletion in HeLa cells following transfection with siSFPQ #1 or siSFPQ #2. *SFPQ* expression level is calculated relative to *GAPDH* and normalized to siCtrl-treated cells. *N* = 4.

E) *Left*, western blot analysis of SFPQ protein abundance in HeLa cells following a modest reduction in *SFPQ* expression upon transfection with siSFPQ #1 or siSFPQ #2. VINCULIN served as a loading control. *Right*, quantification of SFPQ abundance in reference to VINCULIN and normalized to siCtrl-treated cells. *N* = 3.

F) RT-qPCR quantification of the *MBNL1-IPA* and canonical *MBNL1* (e1-e2) transcript levels following modest *SFPQ* depletion. Gene expression level is calculated relative to *GAPDH* and normalized to siCtrl-treated cells. *N* = 4.

G) *MBNL1* pre-mRNA accumulation across the locus following *SFPQ* depletion (with siSFPQ #1). Transcript abundance was measured by RT-qPCR using amplicons spanning e1–i1, ALE–i1, i1–e2 and i9–e10 regions, in reference to *GAPDH* and normalized to siCtrl-treated cells. *N* = 4.

H) Representative western blot analysis of FLAG-SFPQ overexpression in HeLa cells. Cells were transfected with FLAG-SFPQ or a Flag-containing control plasmid. FLAG-SFPQ was detected using an α-FLAG antibody and VINCULIN served as a loading control. *N* = 3.

I) RT-qPCR quantification of the *MBNL1-IPA* and canonical *MBNL1* transcript levels following FLAG-SFPQ overexpression. Gene expression level is calculated relative to *GAPDH* and normalized to siCtrl-treated cells. *N* = 6-7.

#### Figure S5

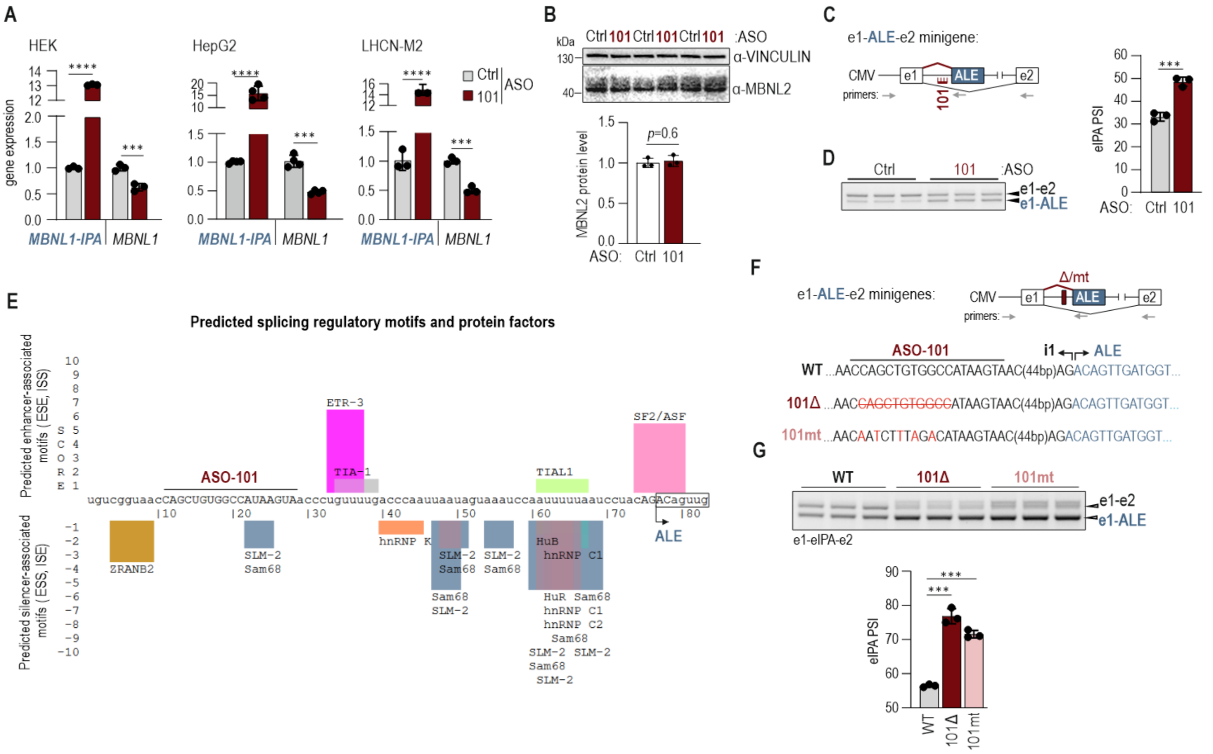


**Supplementary Figure S5. A potent intronic cis-regulatory element represses the *MBNL1* IPA pathway.**

A) RT-qPCR quantification of *MBNL1-IPA* and canonical *MBNL1* transcript levels following treatment with ASO-101 in HEK293, HepG2 and LHCN-M2 cells. Gene expression level is calculated relative to *GAPDH* and normalized to ASO-Ctrl-treated cells. *N* = 3-4.

B) *Top*, western blot analysis of MBNL2 protein abundance following treatment of HeLa cells with ASO-101. VINCULIN served as a loading control. *Bottom*, quantification of MBNL2 protein abundance in reference to VINCULIN and normalized to ASO-Ctrl-treated cells. *N* = 3.

C) Schematic representation of the e1–ALE-e2 minigene reporter showing the ASO-101 target region. Canonical exons, ALE, the ASO-101 target region, and primer positions used for transcript analysis are indicated.

D) *Left*, RT-PCR analysis of transcripts generated from the e1-ALE-e2 minigene in ASO-Ctrl and ASO-101-treated HeLa cells. Products corresponding to the canonical (e1-e2) and ALE-containing (e1-ALE) isoforms are indicated. *Right*, quantification of ALE inclusion is shown as PSI values. *N* = 3.

E) Prediction of RNA regulatory motifs within and around the ASO-101-targeted region using SpliceAid[3]. Predicted enhancer and silencer protein factors are indicated. The position of the ASO-101 target sequence is highlighted, along with the start of ALE. ESE, exonic splicing enhancer; ISS, intronic splicing silencer; ESS, exonic splicing silencer; ISE, intronic splicing enhancer.

F) Schematic representation of wild-type (WT) and mutant e1–ALE-e2 minigene reporters. Nucleotides deleted from the minigene are crossed-out and marked in red (101Δ), nucleotide substitutions introduced into the ASO-101 target region (101mt) are highlighted in red. Canonical exons, ALE, the ASO-101 target region, and primer positions used for transcript analysis are indicated.

G) *Top*, RT–PCR analysis of transcripts generated from WT and mutant e1–ALE-e2 minigene reporters. PCR products corresponding to canonical (e1-e2) and ALE-containing transcripts are indicated. *Right*, quantification of ALE inclusion is shown as PSI values. *N* = 3.

**Figure S6**

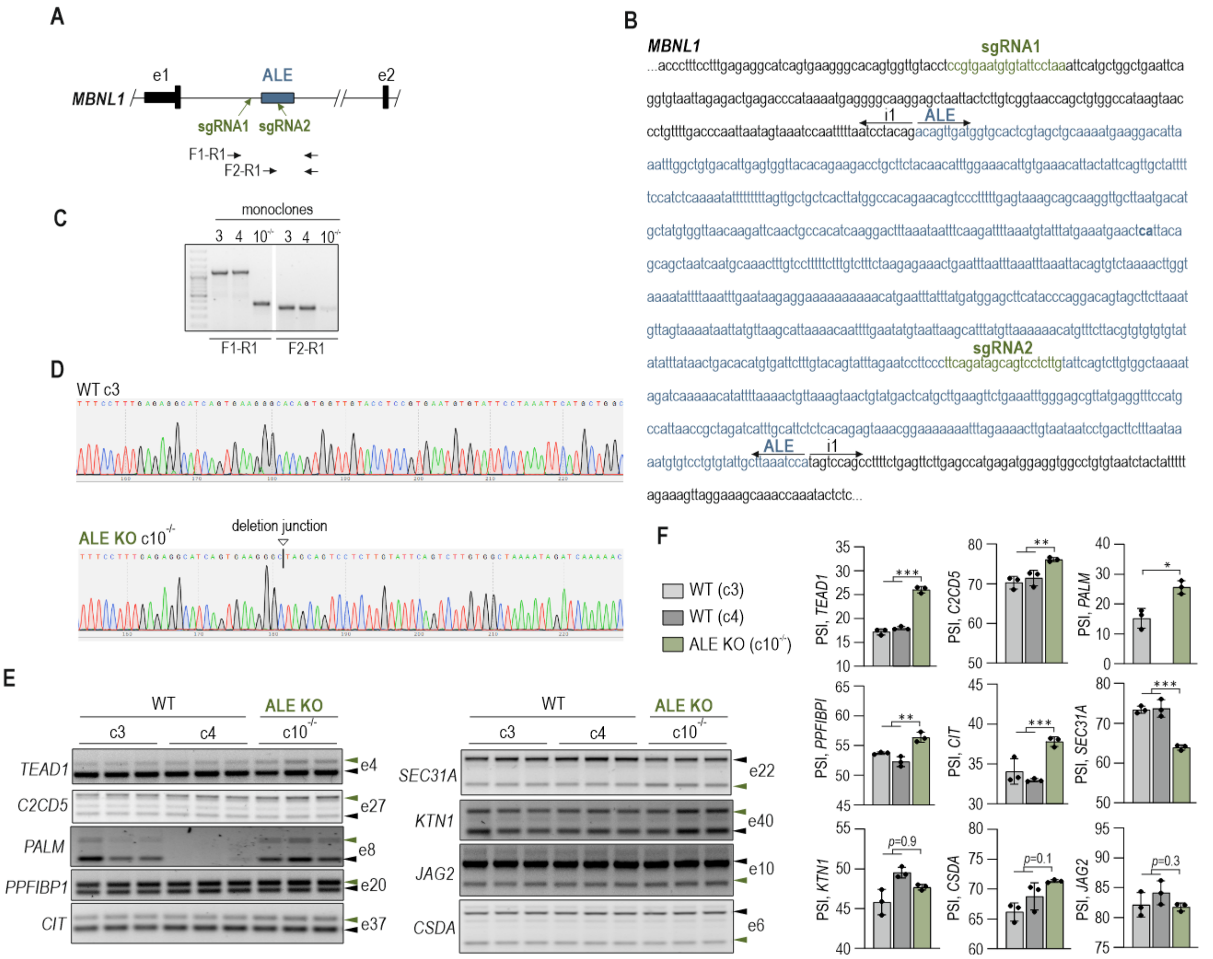


**Supplementary Figure S6. Deletion of the ALE region increases *MBNL1* expression and enhances the MBNL-dependent splicing pattern.**

A) Schematic representation of the *MBNL1* exon 1-ALE region showing approximate positions of sgRNAs used for CRISPR-Cas9-mediated deletion of ALE-associated region (ALE KO). Primers used for genotyping PCR analyses (F1-R1 and F2-R1) are indicated.

B) Genomic nucleotide sequence of the targeted ALE region. ALE sequence is shown in blue and intronic sequences in black. The positions of sgRNA1 and sgRNA2 target sites are marked in green.

C) PCR-based screening of individual HEK293 monoclonal cell lines following CRISPR-Cas9-mediated editing. Primer pairs F1-R1 and F2-R1 were used to identify and select a homozygous ($c10^{-/-}$) ALE KO clone.

D) Representative Sanger sequencing chromatograms confirming CRISPR-Cas9-mediated deletion of the ALE-associated region. Vertical line indicates a breakpoint junction formed after excision of the intervening genomic sequence and repair-mediated ligation of the flanking DNA ends.

E) RT-PCR analysis of MBNL-dependent alternative splicing events in the control (c3, c4) and ALE KO clone ($c10^{-/-}$). Representative analyses of *TEAD* e4, *C2CD5* e27, *PALM* e8, *PPFIBP1* e20, *CIT* e37, *SEC31A* e22, *KTN1* e40, *JAG2* e10 and *CSDA* e6 are shown together with PSI quantification. *KTN1* e40, *JAG2* e10 and *CSDA* e6 events exhibited no statistically significant changes between control and ALE KO cells. Green arrowheads indicate isoforms promoted by MBNL. Statistical significance was determined relative to WT clones, c3 and c4. *N* = 3.

**Figure S7**

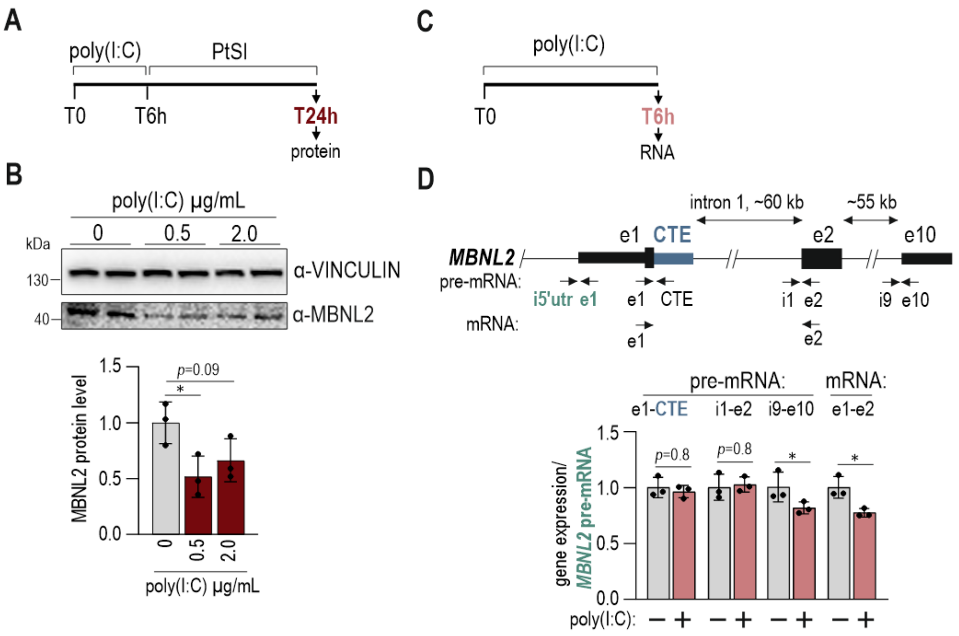


**Supplementary Figure S7. Poly(I:C)-induced innate immune stress remodels *MBNL2* transcript output and reduces MBNL2 protein abundance.**

A) Experimental design of the recovery assay (post-stress induction, PtSI). HeLa cells were treated with poly(I:C) for 6 h and subsequently cultured in fresh medium for additional 18 h before collection for protein analysis.

B) *Left*, representative western blot analysis of MBNL2 protein abundance following 0.5 µg/mL poly(I:C) treatment in HeLa cells. VINCULIN served as a loading control. *Right*, quantification of MBNL2 protein level relative to VINCULIN and normalized to RNAiMAX treated control cells. *N* = 3.

C) Experimental design of poly(I:C) treatment in HeLa cells. Cells were exposed to 0.5 µg/mL poly(I:C) for 6 h and collected for RNA analysis.

D) *Top*, schematic representation of the *MBNL2* locus showing the positions of RT–qPCR primers used to monitor changes in the *MBNL2* pre-mRNA level using primers located near 5'end, across intron 1 and 3'end as well as the mRNA level. Exons are shown as black boxes, a putative *MBNL2* IPA-associated region is shown as a blue box and introns as lines. Distance between selected exons is indicated. *Bottom*, RT-qPCR quantification of *MBNL2* pre-mRNA and mRNA levels in HeLa cells after 6 h treatment with 0.5 µg/mL of poly(I:C). Transcript level was calculated in reference to the internal control, the 5'end of *MBNL2* pre-mRNA which is common for all *MBNL2* isoforms. Such calculation shows changes in transcription and RNA processing in relation to transcriptional activity of the locus in control conditions and upon poly(I:C) treatment. The results were normalized to RNAiMAX treated cells. *N* = 3.

**Figure S8**

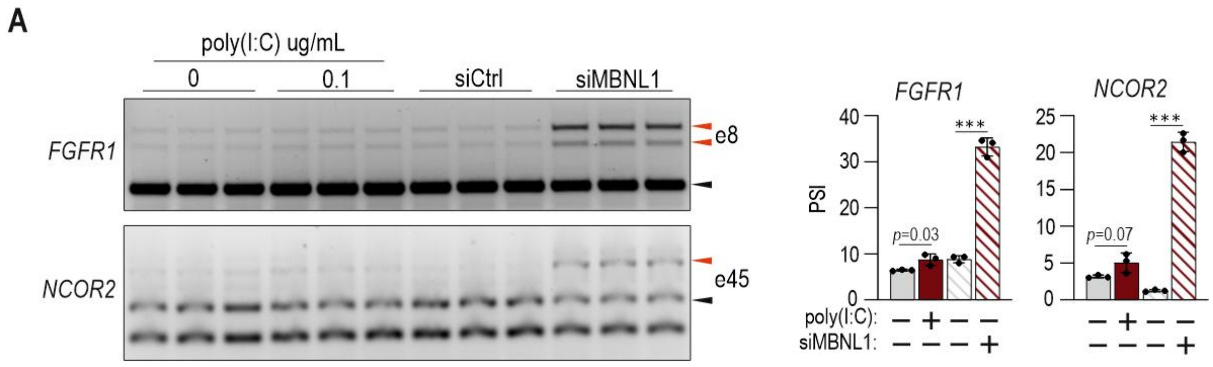


**Supplementary Figure S8. Poly(I:C)-induced innate immune stress reduces MBNL1 abundance and impairs MBNL-dependent splicing in DM1-affected fibroblasts.**

A) *Left*, RT–PCR analysis of selected MBNL-dependent splicing events *NCOR2* e45 and *FGFR1* e8 in DM1-affected fibroblasts following poly(I:C) treatment and PtSI period or siRNA-mediated knockdown of *MBNL1*. *Right*, quantification of alternative splicing events presented as PSI values. *N* = 3.

### SUPPLEMENTARY MATERIALS AND METHODS

#### Genetic constructs

The pEGFP-MBNL1-41 expression vector has been previously described[4]. Flag-SFPQ expression vector has been obtained from Addgene (#166960). The *MBNL1-IPA* CDS expression construct was generated in two successive cloning steps. First, the intervening intronic sequence between exon 1 and the ALE was removed from the e1-ALE plasmid by PCR amplification using primers E1ALE_OE_F and E1ALE_OE_R, followed by plasmid recircularization. Second, the sequence extending from the predicted *MBNL1-IPA* termination codon to the bovine growth hormone polyadenylation signal (bGH-polyA) was removed using primers ALECDS_R and bGH_clon_ALE_F, followed by recircularization of the resulting construct.

#### Cell culture and transfection

HeLa cells (ATCC) were grown in high-glucose DMEM (Lonza) supplemented with 10% fetal bovine serum (FBS) (Sigma Aldrich) and penicillin–streptomycin (Sigma Aldrich). Prior to transfection, the cells were plated in 12-well plates and transfected at 50–60% confluence with siRNA using Lipofectamine RNAiMAX (Thermo Fisher Scientific) according to the manufacturer's protocol. Genes were knocked down with a single siRNA against SFPQ (s12712, Ambion) and a pool of 3 siRNAs (sc-38304, Santa Cruz) and control siRNA (Sigma Aldrich) at 5 nM. The cells were harvested 48 h after transfection.

#### Nanopore data analyses

Publicly available long-read cDNA data from the Jurkat human T-cell leukemia cell line were analysed. The data were released by Oxford Nanopore Technologies as part of the EPI2ME Open Data project (dataset sc_gemx_2025.02; s3://ont-open-data/sc_gemx_2025.02). The dataset comprises two biological replicates. The basecalled BAM files provided by the data providers were used. Reads were converted to FASTQ with samtools (v1.22.1) and aligned to the human reference genome GRCh38.p14 (GCF_000001405.40) using minimap2 (v2.30) in spliced alignment mode. Cell barcodes and UMIs were not extracted or used; reads from all cells and replicates were treated as a single bulk-like pool. Alignments at the *MBNL1-IPA* locus were inspected in IGV (v2.19.6).

#### Sucrose gradient fractionation

Polysome fractionation was performed as previously described[5] with minor modifications. Briefly, approximately 3.5-4 x $10^6$ HEK293 cells per sample were treated with cycloheximide (100 µg/mL; Merck) for 30 min prior to harvesting. Cells were lysed in polysome lysis buffer containing 20 mM Tris-HCl (pH 7.5), 10 mM NaCl, 3 mM $MgCl_2$, 1 mM DTT, 0.3% Triton X-100, 50 mM sucrose, 1 U/µL RNasin (Promega), and 100 µg/mL cycloheximide. Lysates were clarified by centrifugation and loaded onto 15-45% linear sucrose gradients. Gradients were centrifuged at 160,000 x *g* for 2 h 40 min at 4°C using an SW41 Ti rotor (Beckman Coulter). Twelve fractions were collected using a Piston Gradient Fractionator (BioComp Instruments) coupled to a 2110 Fraction Collector (Bio-Rad) while monitoring absorbance at 254 nm. RNA was recovered from individual fractions by isopropanol precipitation and purified using TRI Reagent (Merck) according to the manufacturer's instructions. Purified RNA was subsequently used for cDNA synthesis and RT-qPCR analyses. Primer sequences used for RT-qPCR are provided in Supplementary Table S1. The distribution of individual transcripts across the gradient was calculated as previously described[6]. Relative abundance in each fraction was determined using the formula: $\Delta CT = CT_{fraction\ 1} - CT_{fraction\ X}$. Fraction distribution (%) = $(2^{\Delta CT\ fraction\ X} \times 100) / \Sigma(2^{\Delta CT\ fractions\ 1–12})$.

#### Micropeptide structure prediction

The amino acid sequence of the predicted MBNL1-IPA micropeptide was generated by in silico translation of the MBNL1-IPA open reading frame and submitted to AlphaFold2[7] for structure prediction. The highest-ranking model was selected for further analysis. Prediction confidence was evaluated using predicted local distance difference test (pLDDT) scores. Structural visualization and figure preparation were performed using PyMOL v2.5.4.

#### Immunoblotting

Cells were lysed in RIPA buffer (150 mM NaCl, 50 mM Tris-HCl pH 8.0, 1 mM EDTA, 0.5% NP-40, 0.5% Triton X-100, 0.5% sodium deoxycholate, and 0.1% SDS) supplemented with Halt™ EDTA-free Protease Inhibitor Cocktail (Thermo Fisher Scientific). Lysates were incubated on ice for 30 min, snap-frozen at −80°C overnight, and clarified by centrifugation at 18,000 x g for 15 min at 4°C. Protein concentrations were determined using the Pierce BCA Protein Assay Kit (Thermo Fisher Scientific). Equal amounts of protein (10-15 µg) were mixed with Bolt™ reducing agent and Bolt™ LDS sample buffer (Thermo Fisher Scientific), incubated at 70°C for 10 min, and separated on Bolt™ 4–12% Bis-Tris Plus gradient gels (Thermo Fisher Scientific). Proteins were transferred onto 0.1-µm nitrocellulose membranes (Merck) for micropeptide detection. Wet transfer was performed at 100 V for 1 h at 4°C. Membranes were blocked for 1 h in 5% (w/v) skim milk prepared in TBST (Tris-buffered saline supplemented with 0.1% Tween-20) and incubated with primary antibodies listed in Supplementary Table S1. HRP-conjugated secondary antibodies were used where appropriate. Immunoreactive signals were detected using Immobilon™ Forte Western HRP substrate (Sigma-Aldrich) and visualized using a ChemiDoc™ imaging system (Bio-Rad).

**Supplementary Table S1 (Excell).** List of ASOs, siRNAs, primers and antibodies.
